# Machine learning prediction of cancer cell metabolism from autofluorescence lifetime images

**DOI:** 10.1101/2022.12.16.520759

**Authors:** Linghao Hu, Nianchao Wang, Joshua D Bryant, Lin Liu, Linglin Xie, A Phillip West, Alex J. Walsh

**Affiliations:** Department of Biomedical Engineering, Texas A&M University, College Station, TX 77843; Microbial Pathogenesis& Immunology, Health Science Center, Texas A&M University, College Station, TX, 77807; Department of Nutrition, Texas A&M University, College Station, TX, 77030

**Keywords:** fluorescence lifetime, metabolism, machine learning, convolutional neural network

## Abstract

Metabolic reprogramming at a cellular level contributes to many diseases including cancer, yet few assays are capable of measuring metabolic pathway usage by individual cells within living samples. Here, we combine autofluorescence lifetime imaging with single-cell segmentation and machine-learning models to predict the metabolic pathway usage of cancer cells. The metabolic activities of MCF7 breast cancer cells and HepG2 liver cancer cells were controlled by growing the cells in culture media with specific substrates and metabolic inhibitors. Fluorescence lifetime images of two endogenous metabolic coenzymes, reduced nicotinamide adenine dinucleotide (NADH) and oxidized flavin adenine dinucleotide (FAD), were acquired by a multi-photon fluorescence lifetime microscope and analyzed at the cellular level. Quantitative changes of NADH and FAD lifetime components were observed for cells using glycolysis, oxidative phosphorylation, and glutaminolysis. Conventional machine learning models trained with the autofluorescence features classified cells as dependent on glycolytic or oxidative metabolism with 90 – 92% accuracy. Furthermore, adapting convolutional neural networks to predict cancer cell metabolic perturbations from the autofluorescence lifetime images provided improved performance, 95% accuracy, over traditional models trained via extracted features. In summary, autofluorescence lifetime imaging combined with machine learning models can detect metabolic perturbations between glycolysis and oxidative metabolism of living samples at a cellular level, providing a label-free technology to study cellular metabolism and metabolic heterogeneity.

## Introduction

Cellular metabolism is linked with cellular function for many cell types. For example, cancer is often characterized by a dependence on aerobic glycolysis and immune cell function is dependent on glycolysis (1, 2). Shifts in cellular metabolic pathways can be dynamic and heterogeneous across tissues in response to microenvironments, manipulations, and cell functions. Single-cell RNA analyses have revealed that metabolic heterogeneity affects cancer metastasis and functions of T cells (3, 4). However, a full understanding of the spatial and dynamic nature of metabolic heterogeneity on cancer development, invasion, and drug response is inaccessible because we lack technologies that can measure the metabolism of living samples with cellular resolution.

Currently, metabolic measurement technologies are limited in spatial and temporal resolution. The oxygen consumption rate (OCR) and the extracellular acidification rate (ECAR) of cell populations can be used to evaluate mitochondrial respiration and glycolysis, respectively (5, 6). However, these measurements are recorded for cell populations. Likewise, biochemical analyses of metabolic enzymes, including Western Blot analysis, mass spectroscopy, mRNA analysis, and immunohistochemistry, require cell or tissue fixation and generally lack single-cell resolution (7). Positron Emission tomography (PET) detects radioactive substances, such as 2-deoxy-2-[18F] fluoro-D-glucose (FDG) to visualize glucose uptake of tumors which is higher than surrounding tissue due to enhanced glycolysis (8). FDG-PET is used clinically, and additional contrast agents are in development to detect choline metabolism, glutamine transport, and fatty acid metabolism (9–11). However, the spatial resolution of PET is fundamentally limited by the millimeter distances that positrons can travel prior to annihilation events (12). Signaling pathways can be labeled with genetically-encoded fluorescence proteins to offer metabolic information, however, the dependence on exogenous fluorescent labels requires cellular manipulations and brings confounding factors like cell destruction, dye concentration, and label distribution errors (13, 14). Raman spectroscopy can observe the redox state of cytochrome and mitochondrial dysfunction, but its usage is limited by low sensitivity and resolution (15–17). Therefore, a robust technology capable of live-cell metabolic measurements remains elusive, yet potentially impactful due to the large number of diseases and pathologies characterized by metabolic dysfunction and heterogeneity.

Autofluorescence imaging of two key endogenous metabolic co-enzymes reduced nicotinamide adenine dinucleotide (NADH) and oxidized flavin adenine dinucleotide (FAD) offers functional metrics for detecting metabolic variations (18, 19). NADH and FAD are used in metabolic pathways including glycolysis, oxidative phosphorylation (OXPHOS), and glutaminolysis. The fluorescence intensity ratio of NADH and FAD is defined as the optical redox ratio and has been widely used as a marker of the redox state in cells and tissues (19, 20). The spectral properties of NADH and its phosphorylated form, NADPH are identical, thus NAD(P)H is used to represent their combined fluorescence signal detected from cells. The fluorescence lifetime is the time a fluorophore remains in the excited state before releasing a fluorescent photon and resolves information on chemical structures and the surrounding microenvironments of NAD(P)H and FAD (21–23). Both NAD(P)H and FAD can exist in two conformations, protein-bound or free within cells, each of which has a different fluorescence lifetime (22, 23). Fluorescence lifetime imaging (FLIM) resolves the fraction of free and protein-bound coenzymes as well as the corresponding short- and long-lifetime components (18, 24). The fluorescence intensity and lifetime of NAD(P)H and FAD are sensitive to metabolic changes in precancerous tissues, disease development, drug treatment responses of cancer cells, differentiation of stem cells, and macrophage phenotype (20, 25–34). Moreover, cell segmentation from the NAD(P)H intensity images provides single-cell and subcellular information and quantifies cellular heterogeneity (35, 36).

Although autofluorescence imaging of NAD(P)H and FAD often detects metabolic perturbations between samples, the lifetime metrics lack specificity for metabolic pathways. In this paper, we define the effects of metabolic pathway perturbations on NAD(P)H and FAD fluorescence lifetime metrics and create and validate machine learning (ML) models that predict glycolytic or oxidative phenotypes of cells from autofluorescence lifetime features. First, alterations in NAD(P)H and FAD fluorescence lifetime features are defined for metabolic pathway perturbations of cancer cells using both chemical inhibition and substrate manipulation of glycolysis, oxidative phosphorylation, and glutaminolysis. Then, we compare extracted feature-based machine learning models with convolutional neural networks (CNNs) for the prediction of cellular metabolic states from autofluorescence lifetime images. The models are then evaluated across different metabolic perturbations and cell types to evaluate model robustness and transference. The results and metabolism-prediction ML models address current limitations in the interpretation of autofluorescence lifetime datasets and promote the application of autofluorescence imaging for live cell studies of cancer cell metabolism and metabolically-targeted drug studies.

## Materials and Methods

### Cell culture and preparation

MCF7 breast cancer cells were cultured in high glucose Dulbecco’s Modified Eagle’s Medium (DMEM), supplemented with 1% antibiotic-antimycotic, and 10% fetal bovine serum (FBS). For fluorescence lifetime imaging, cells were seeded at a density of 2 × 10^5^ per 35mm glass-bottom imaging dish 48 hours before imaging. Each dish was refreshed with the culture media 30 minutes before imaging to ensure constant consistent nutrient concentrations while imaging. For glucose and OXPHOS inhibition experiments, two types of culture media were prepared for different treatments: control media 1 (CM1): DMEM (Gibco™, 10313021) with high glucose (25 mM) and pyruvate (1 mM), but without glutamine; and control media 2 (CM2) glucose starvation media: DMEM (Gibco™, A1443001) without glucose, glutamine, or pyruvate. Different concentrations of 2-Dexoy-D-glucose (10 mM, 20 mM, 50 mM, Thermo Scientific, AC111980050) were added to cells in the CM1 1 hour before imaging to allow reactions to inhibit the glycolysis pathways (39). For a glucose depletion group, MCF7 cells were seeded on imaging dishes in CM1 as described and the media was exchanged for CM2 supplemented with pyruvate (50 mM, Gibco™, 11360070) 1 hour before imaging. For the sodium cyanide (NaCN) group, cells were seeded in CM1 and NaCN (4 mM, SIGMA-ALDRICH, 380970) was added to CM1 to inhibit OXPHOS 5 minutes before imaging (25). In the pyruvate concentration imaging experiment, CM2 was supplemented with titrated concentrations of pyruvate (0 mM, 10 mM, 20 mM, 50 mM). Cells were exposed to the media with different concentrations of pyruvate for 1 hour before imaging. For targeting glutaminolysis, three types of culture media were prepared for different treatments: control media 3 (CM3): DMEM with glucose (25 mM) and pyruvate (1 mM), and glutamine (2 mM, Gibco™, 25030081), glutamine depletion media: DMEM with glucose (25 mM) and pyruvate (1 mM), but without glutamine, and glutamine-only media: DMEM with glutamine (2 mM), but without glucose or pyruvate. To inhibit glutaminolysis, bis-2-(5-phenylacetamido-1,3,4-thiadiazol-2-yl) ethyl sulfide (BPTES, 10 μm, ASTATECH, A11656) was added to the cells in CM3 1 hour before imaging (35). To isolate the glutaminolysis pathway over time, cells were plated in CM3 and the media was exchanged for the glutamine-only media, and the cells were imaged at 1, 2, and 3 hours. The concentration of metabolic substrates in each group is summarized in the Sup. Table 1.

HepG2 hepatoma cells were cultured in low glucose (5.6 mM) DMEM supplemented with 1% penicillin-streptomycin, and 10% FBS. The cells were seeded at a density of 2 × 10^5^ per 35 mm glass-bottom imaging dish and fasted for 24 hours. Two metabolic groups were prepared by treating the cells with either 30 mM glucose or 0.4 mM palmitate (PA) respectively 12 hours before imaging. The NAD(P)H and FAD fluorescence lifetime images from the activated and quiescent T cells were provided by AJ Walsh and MC Skala (35).

### Fluorescence lifetime imaging and analysis

The NAD(P)H and FAD fluorescence lifetime images were obtained using a custom-built multi-photon microscope (Marianas, 3i) coupled with a 40X water immersion objective (1.1 NA) and a tunable (680nm – 1080nm) Ti: sapphire femtosecond laser (COHERENT, Chameleon Ultra II). All cell imaging dishes were placed in a stage-top incubator (okolab) to maintain the environment of the cells at 37 °C, 5% CO_2_, and 85% relative humidity while imaging. NAD(P)H fluorescence was excited at 750 nm with a laser power of 18 mW to 20 mW. FAD fluorescence was excited at 890 nm with a laser power of 25 mW to 28 mW. To isolate NAD(P)H and FAD fluorescence emission, a 447/60 nm bandpass filter and a 550/88 nm bandpass filter, respectively, were placed before each detector. Fluorescence lifetime images of NAD(P)H and FAD were obtained sequentially by photomultiplier tube (PMT) detectors (HAMAMATSU) attached to a time-correlated single-photon counting (TCSPC) electronics module (SPC-150N, Becker & Hickl). Each fluorescence lifetime image (256 × 256 pixels, 270μm x 270μm) was acquired with a pixel dwell time of 50 μs and 5 frame repeats for a collection time of 60 seconds. Both NAD(P)H and FAD fluorescence lifetime images were captured in at least five randomly selected positions for each dish, and three technical replicates were performed to ensure the reliability of the results. The second harmonic generated signal of urea crystals was excited at 900 nm and measured with the NAD(P)H channel for the instrument response function (IRF). Fluorescence lifetime measurements of the system were validated with a YG fluorescent bead, which had a measured lifetime of 2.1 ns, consistent with previously published values (32).

### Cell-based image analysis

Fluorescence lifetime decays were analyzed by SPCImage (Becker & Hickl). Different thresholds were used to exclude pixels with low fluorescence intensity in NAD(P)H (minimum threshold of peak = 20 photons) and FAD (minimum threshold of peak = 3 photons) fluorescence lifetime images. A binning of nine surrounding pixels was used, and the decay curve of each pixel was deconvoluted from the measured IRF of urea crystals and fitted to a two-component exponential model, *I*(*t*) = *α*_1_*e*^−*t/τ*_1_^ + *α*_2_*e*^−*t/τ*_2_^ + *C*, where *I*(*t*) represents the fluorescence intensity as a function of time *t*, *α_1_*, *α_2_* are the fractions of the short and long fluorescence lifetime, respectively, and their sum is 100 percent (*α*_1_ + *α*_2_ = 1). *τ_1_*, *τ_2_* are the corresponding short and long lifetimes, and *C* accounts for background light. NAD(P)H has a short lifetime at the free conformation, and a long lifetime when bound to a protein. Conversely, FAD has a long lifetime when it is free, and a short lifetime when bound.

Images were then segmented into individual cell, cytoplasm, and nucleus compartments to acquire cell-based fluorescence lifetime endpoints. The cell segmentation process was based on the NAD(P)H intensity images and achieved in CellProlifer using a customized pipeline. Firstly, the raw NAD(P)H intensity images were rescaled between 0 and 1. Next, the backgrounds were extracted by using default object identification in CellProlifer. Since the nuclei in the cells were darker compared with surrounding cytoplasmic regions, the image features of speckles and dark holes were enhanced to improve the appearance of nucleus regions. Then an adaptive Otsu method was applied to identify the nucleus within a typical diameter range (5~20 for MCF7 cells), and the diameter can be adjustable for different cells. To distinguish between individual nuclei that are touching each other, each object was identified by the peak of brightness and the dividing line was determined by the occurrence of indentation. Then, the cellular regions were identified by propagating from the pre-identified nucleus regions through an adaptive Otsu method. The threshold correction factor and size of the adaptive window were adjusted to optimize the performance of the Otsu method in different cells. Cytoplasm masks were determined by subtracting the nucleus objects from cell objects. Finally, all identified cells were filtered based on the area of the cytoplasm to exclude clumped cells and error objects. The cell number of each group is summarized in Table 1 and Sup. Table 2.

**Table 1.**
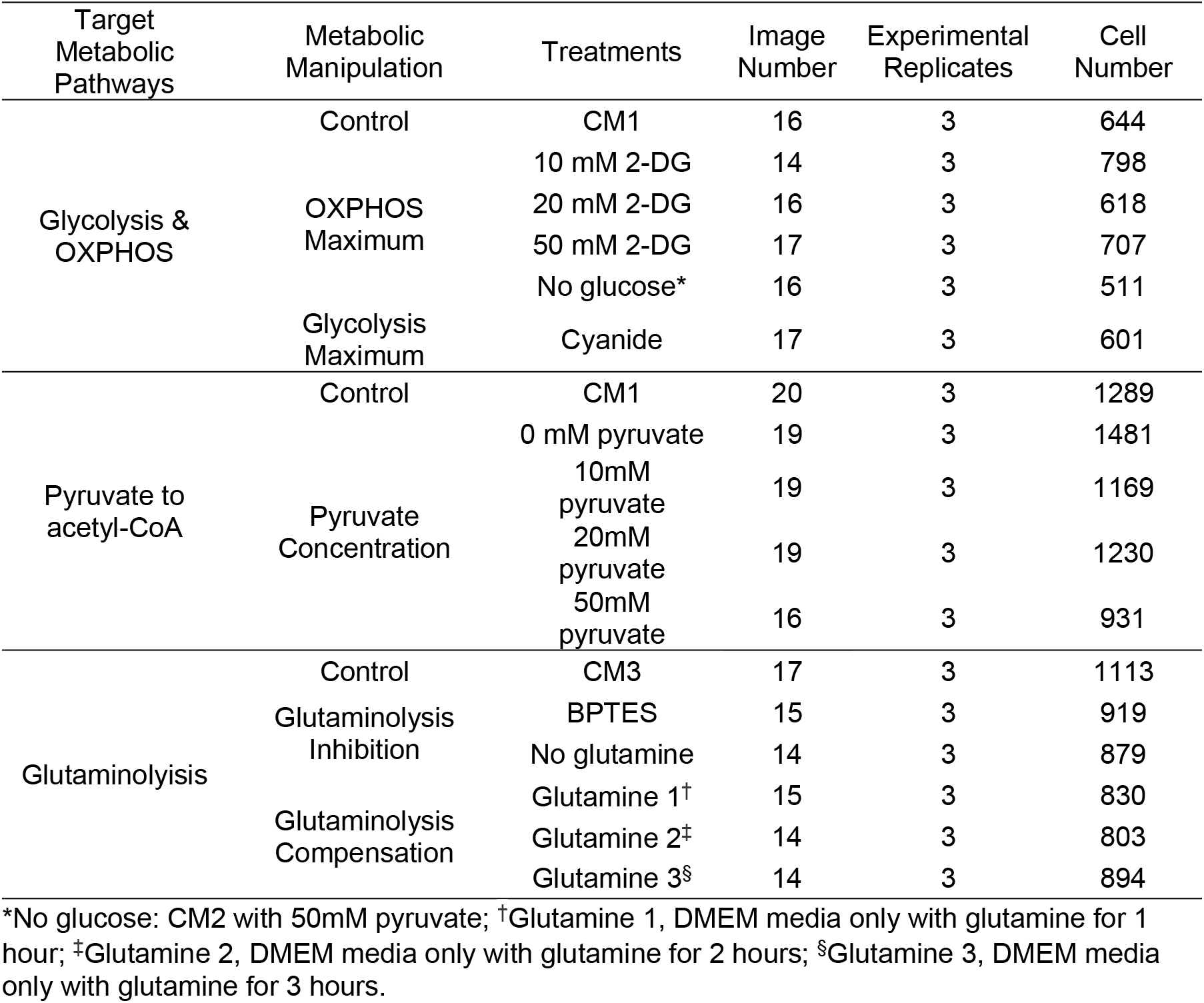
Number of Cells in each group

Image processing was performed using MATLAB to calculate the images of optical redox ratio (FAD fluorescence intensity divided by the summed intensity of FAD and NAD(P)H, FAD/(NAD(P)H + FAD)), weighted average fluorescence lifetime (*τ_m_* = *α*_1_*τ*_1_ + *α*_2_*τ*_2_), and fluorescence lifetime redox ratio (FLIRR, bound NAD(P)H fraction divided by the bound FAD fraction, NAD(P)H *α_2_*/FAD *α_1_*). Twelve NAD(P)H and FAD fluorescence features including optical redox ratio, FLIRR, NAD(P)H *τ_m_*, NAD(P)H *τ_1_*, NAD(P)H *τ_2_*, NAD(P)H *α_1_*, NAD(P)H intensity, FAD *τ_m_*, FAD *τ_1_*, FDA *τ_2_*, FAD *α_1_*, and FAD intensity were averaged across all pixels within a cytoplasm for each segmented cell for cell-level data.

### Statistical analysis and classification

Data analysis was performed in R Studio. A two-sided student t-test with a Bonferroni correction was used to indicate differences across cell groups for each fluorescence lifetime endpoint, and an alpha significance value of 0.05 was used to indicate significance. The Uniform Manifold Approximate and Projection (UMAP) method was used to visualize clustering within the autofluorescence imaging datasets (76). Cancer cells treated with sodium cyanide were defined as the OXPHOS inhibition group. Cells treated with 50 mM 2-DG and exposed to no glucose media were identified as the glycolysis inhibition group. Classical machine learning algorithms (random forest tree (RFT), support vector machine (SVM), quadratic discriminant analysis (QDA)) were trained to classify cells with inhibited glycolysis versus cells with inhibited OXPHOS based on the fluorescence lifetime features. These models were trained on a randomly selected 75% (1365 cells) of the dataset and tested on the remaining 25% (454 cells) of the dataset. A receiver operation characteristic (ROC) curve and confusion matrix were used to evaluate the performance of the models on test datasets. The importance within the random forest tree classification model and the AUC (area under the curve) value of the ROC curve of each fluorescence lifetime feature were used to assess each feature’s contribution to the prediction. Each model was tested with 5-fold cross-validation and an average accuracy was computed to ensure its robustness. When predicting the metabolic activities of liver cancer cells, to avoid lifetime parameter differences within cell types, each feature value was normalized with the mean value of corresponding control cells to get the relative changes, and a new model was trained with normalized FLIM features.

### Image preprocessing for CNN

Since CNN development requires a larger dataset than the classical machine learning algorithms, the glycolysis and OXPHOS inhibition experiments were repeated to obtain ~5000 original cells (Sup. Table 3). Each cell was extracted based on the bounding box of its mask generated by CellProfiler to produce six autofluorescence lifetime endpoint images (NAD(P)H *τ_1_*, NAD(P)H *τ_2_*, NAD(P)H *α_1_*, NAD(P)H *τ_m_*, NAD(P)H intensity, and FAD intensity). Then, the following image preprocessing procedure (Figure 1) was achieved in python running on Jupyter Notebook on the platform of Anaconda3 with the help of the OpenCV package. To remove incomplete or uninformative cells, the cells were filtered by thresholding the entropy of NAD(P)H intensity and NAD(P)H *τ_m_* images, and the threshold values were defined according to the distribution of entropy with a Gaussian approximation. Since the CNN classifier requires inputs of uniform size, we padded all cancer cells to be 40 × 40 pixels with black borders. The padding size was decided by the largest cell size and the cell size distribution of the dataset. After noise removal and padding, a montage of images of the cells in each metabolic group was visually inspected. Finally, the original training dataset was augmented by rotating each cell image by 45, 90, 135, 180, and 270 degrees, and also by flipping the cell image horizontally and vertically. This procedure amplified the size of the original training dataset by seven times (Sup. Table 3). To test the metabolic prediction performance of different autofluorescence lifetime features by CNN, input datasets consisted of all NAD(P)H fluorescence lifetime components (NADH *τ_1_*, NADH *τ_2_*, NADH *α_1_*, NADH *τ_m_*, NADH intensity, FAD intensity) or a subset of these images (Figure 1).

**Figure 1.**
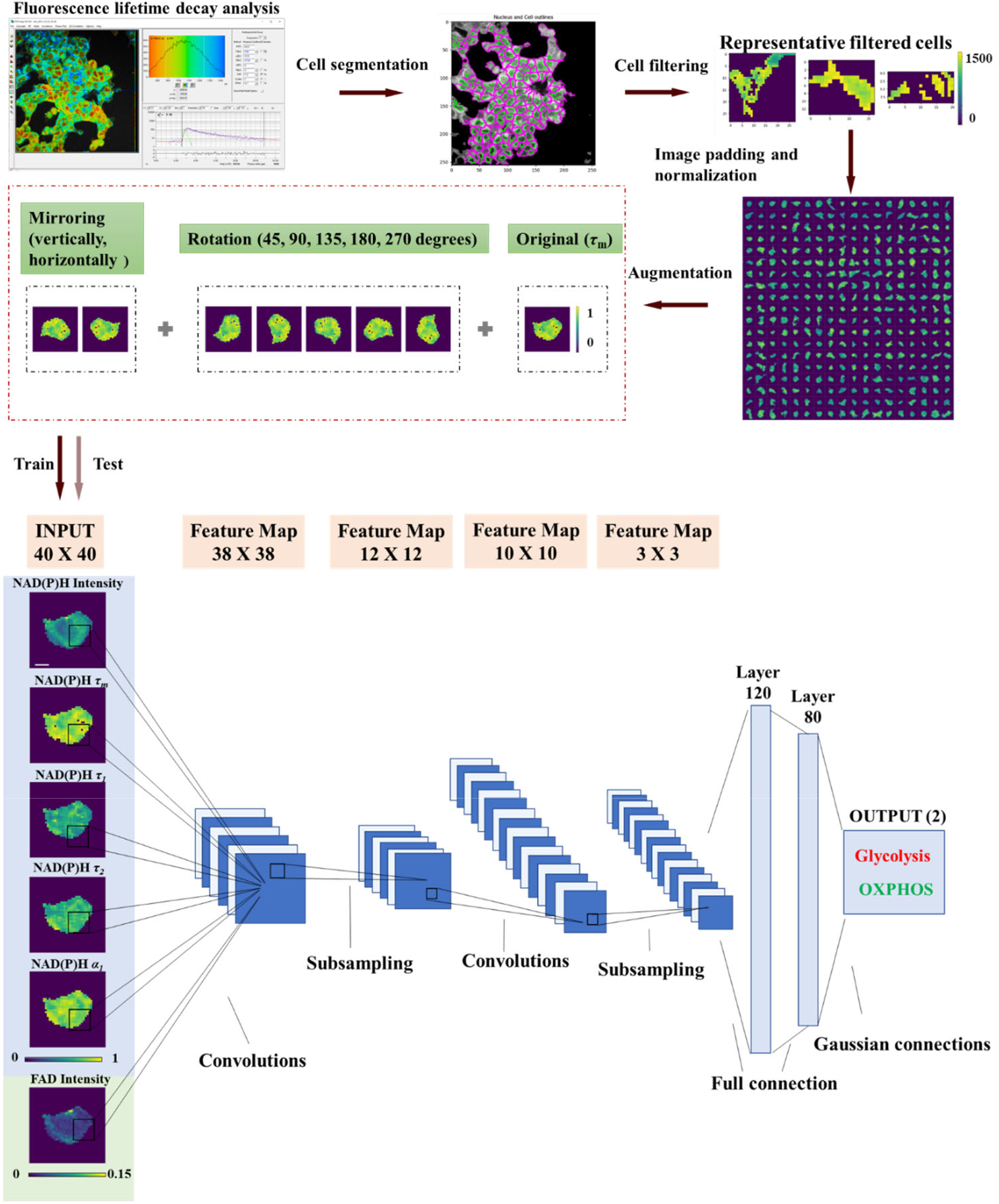
Cancer cell image data preprocessing, and CNN development workflow. Fluorescence lifetime decay was analyzed by SPCImage to extract different endpoints (intensity, *α_1_*, *τ_1_*, *τ_2_*), and the cells were then segmented by CellProfiler from NAD(P)H intensity images. The images were padded to be 40 × 40, and augmented by rotating different angles (45, 90, 135, 180, 270) and mirroring. Finally, LeNet CNN models were trained with different fluorescence lifetime component images (NADH *τ_1_*, NADH *τ_2_*, NADH *α_1_*, NADH *τ_m_*, NADH intensity, FAD intensity), scale bar = 7 μm.

### CNN development

A LeNet architecture for the CNN was developed using the machine-learning library Keras with a Tensorflow backend in python running on Jupyter Notebook on the platform Anaconda 3. The model was composed of two convolutional layers and two pooling layers (77). The input layer was adjusted to 40 × 40 with different channel numbers to fit the input number of lifetime components (Figure 1). For the training, 60% of the cells were randomly selected as the training dataset, 10% of cells were defined as the validation dataset, and the remaining 30% were used to test the model (Sup. Table 3). The network was trained using an NVIDIA GeForce RTX 2070 GPU, and the training parameters including learning rate, batch size, and epochs were tuned to achieve the best performance of the model. The AUC, accuracy, precision, recall, precision-recall curve, and ROC curve of the prediction on the test dataset were used to evaluate the performance of CNN models. To compare the performance of CNN with classical machine learning models, we trained three typical machine learning algorithms (random forest tree, support vector machine, and quadratic discriminant analysis) with the same datasets. We calculated the average value of nonzero pixels in each input image and used the average value as a feature to train the classical machine learning models for the classification of glycolysis inhibited or OXPHOS inhibited cells.

### Seahorse metabolic flux assay

A Seahorse XFe96 extracellular flux analyzer (Seahorse Biosciences, Santa Clara, CA) was used to assess the mitochondrial and glycolytic function of the cells in different metabolic groups. MCF7 breast cancer cells were plated at two densities (5 × 10^5^ cells/ml, 10^6^ cells/ml) on a Seahorse 96-well plate in a DMEM-based medium without phenol red, bicarbonate, glucose, pyruvate, or glutamine. Pyruvate (1 mM), glucose (10 mM), 2-DG (50 mM), and cyanide (4 mM) were sequentially injected into the media. Oxygen consumption rate (OCR) and extracellular acidification rate (ECAR) were measured every 5 minutes for 15 cycles.

## Results

### NAD(P)H fluorescence lifetime imaging reveals glycolytic and OXPHOS states of cancer cells

In the autofluorescence images of NAD(P)H of MCF7 cells, the nuclei were darker than the cytoplasm because NAD(P)H was primarily located in the cytosol and mitochondria (Figure 2(a)). Metabolic perturbations to enhance and inhibit glycolysis altered autofluorescence lifetime features averaged across the cytoplasm pixels of each segmented cell (Table 2, Figure 2). Inhibition of glycolysis within MCF7 cells with 2-DG treatment and glucose starvation resulted in a decreased NAD(P)H free fraction (*α_1_*), an increased free NAD(P)H fluorescence lifetime (*τ_1_*), and an increased bound NAD(P)H fluorescence lifetime (*τ_2_*) as compared with control MCF7 cells (Table 2, Figure 2(b)(c)(d)). These NAD(P)H lifetime variations led to a longer NAD(P)H mean lifetime (*τ_m_*) of MCF7 cells with inhibited glycolysis (Figure 2(e)). Additionally, titrated concentrations of 2-DG promoted consistent reductions in NAD(P)H free fraction (*α_1_*) and increases in NAD(P)H fluorescence lifetimes (*τ_1_*, *τ_2_*, *τ_m_*). Conversely, an increased NAD(P)H free fraction (*α_1_*), and shorter free and bound NAD(P)H fluorescence lifetime (*τ_1_*, *τ_2_*) were observed in cyanide-treated MCF7 cells, relative to the corresponding values of control cells (Table 2, Figure 2(b)(c)(d)). These lifetime variations resulted in a shorter NAD(P)H mean lifetime (*τ_m_*) for MCF7 cells with OXPHOS inhibition (Figure 2(e)). With OXPHOS inhibition by cyanide, the NAD(P)H intensity of cancer cells increased, and the FAD intensity decreased, causing a decrease in the intensity redox ratio (IRR, FAD/(FAD+NAD(P)H)) after cyanide exposure (Sup Fig 1, Table 2). Glucose starvation resulted in a significant decrease in both NAD(P)H and FAD intensities, generating a lower intensity redox ratio (Sup Fig 1, Table 2). Glucose starvation altered the FAD fluorescence lifetime features of MCF7 cells resulting in an increased mean FAD lifetime (Table 2, Sup Fig 2).

**Figure 2.**
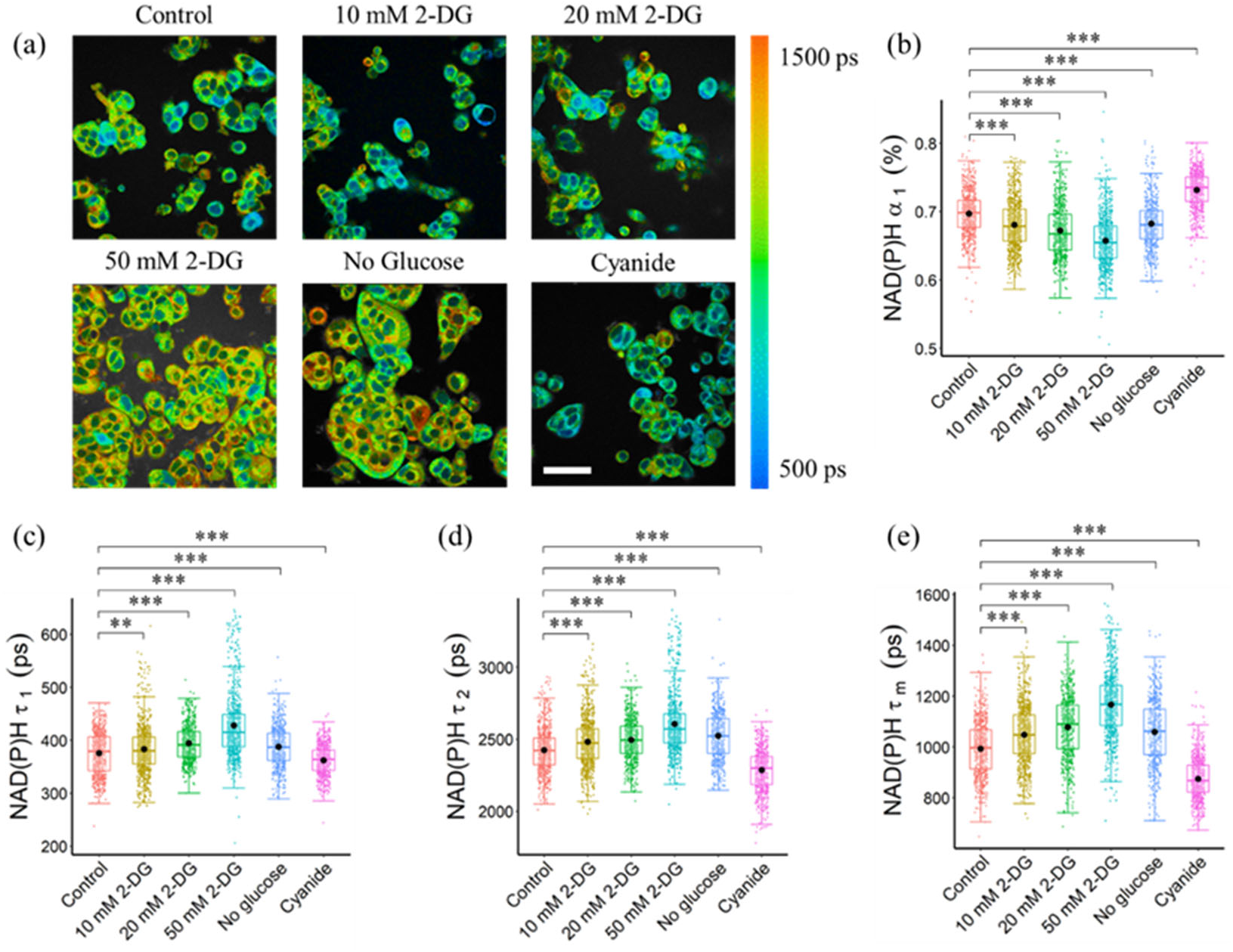
Glycolysis and OXPHOS inhibition altered the NAD(P)H lifetime of MCF7 cells. (a) Representative NAD(P)H *τ_m_* images, scale bar = 60 μm. (b) NAD(P)H *α_1_* (c) NAD(P)H *τ_1_* (d) NAD(P)H *τ_2_* and (e) NAD(P)H *τ_m_* of MCF7 cells exposed to different metabolic environments. ***P < 0.001, **P < 0.01 for two-sided student test with Bonferroni correction for multiple comparisons. Substrates in each media: Control (25 mM glucose + 1 mM pyruvate), 2-DG (25 mM glucose + 1 mM pyruvate + 10/20/50 mM 2-DG), No glucose (50 mM pyruvate), Cyanide (25 mM glucose + 1 mM pyruvate + 4 mM NaCN).

**Table 2.**
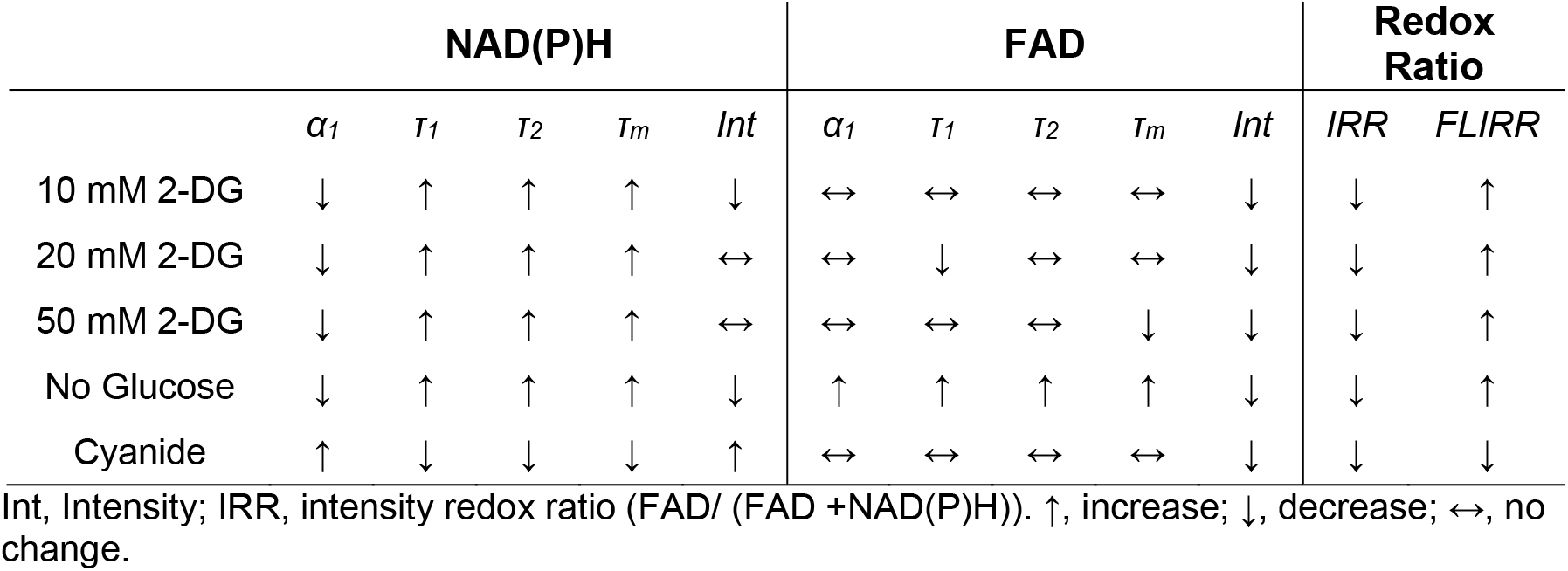
Changes of NAD(P)H and FAD fluorescence lifetime under different metabolic perturbations

|  | NAD(P)H |  |  |  |  | FAD |  |  |  |  | Redox Ratio |  |
| --- | --- | --- | --- | --- | --- | --- | --- | --- | --- | --- | --- | --- |
| | $\alpha_1$ | $\tau_1$ | $\tau_2$ | $\tau_m$ | Int | $\alpha_1$ | $\tau_1$ | $\tau_2$ | $\tau_m$ | Int | IRR | FLIRR |
| 10 mM 2-DG | ↓ | ↑ | ↑ | ↑ | ↓ | ↔ | ↔ | ↔ | ↔ | ↓ | ↓ | ↑ |
| 20 mM 2-DG | ↓ | ↑ | ↑ | ↑ | ↔ | ↔ | ↓ | ↔ | ↔ | ↓ | ↓ | ↑ |
| 50 mM 2-DG | ↓ | ↑ | ↑ | ↑ | ↔ | ↔ | ↔ | ↔ | ↓ | ↓ | ↓ | ↑ |
| No Glucose | ↓ | ↑ | ↑ | ↑ | ↓ | ↑ | ↑ | ↑ | ↑ | ↓ | ↓ | ↑ |
| Cyanide | ↑ | ↓ | ↓ | ↓ | ↑ | ↔ | ↔ | ↔ | ↔ | ↓ | ↓ | ↓ |

Int, Intensity; IRR, intensity redox ratio (FAD/ (FAD +NAD(P)H)). ↑, increase; ↓, decrease; ↔, no change.

The effects of pyruvate and glucose, as well as 2-DG and cyanide treatments, on glycolysis and mitochondrial respiration of MCF7 cells were measured with a Seahorse analyzer. The ECAR increased when glucose was added, and decreased with 2-DG injection, while no changes were observed for the addition of pyruvate or cyanide (Sup Fig 3(a)). The OCR decreased with cyanide injection (Sup Fig 3(b)).

### NAD(P)H fluorescence lifetime imaging reveals glutaminolysis perturbations within cancer cells

The stimulation and inhibition of the glutaminolysis pathway within MCF7 cells altered NAD(P)H fluorescence lifetimes. Inhibition of glutaminolysis with both glutamine starvation and BPTES treatment increased the NAD(P)H free fraction (*α_1_*), and decreased the free and bound NAD(P)H fluorescence lifetimes (*τ_1_*, *τ_2_*) (Figure 3(b)(c)(d)), resulting in a decreased mean NAD(P)H lifetime (*τ_m_*) (Figure 3(e)). Furthermore, an increase in the intensity redox ratio was observed in the BPTES-treated cells compared with the control group (Figure 3(f)). Glutamine starvation increased the bound fraction of FAD (*α_1_*) and lowered the bound and free lifetimes (*τ_1_*, *τ_2_*) resulting in a lower mean FAD lifetime (*τ_m_*) (Sup Fig 4). BPTES treatment of MCF7 cells increased the lifetime of free FAD (*τ_2_*) but did not affect the other FAD lifetime parameters (Sup Fig 4(c)).

**Figure 3.**
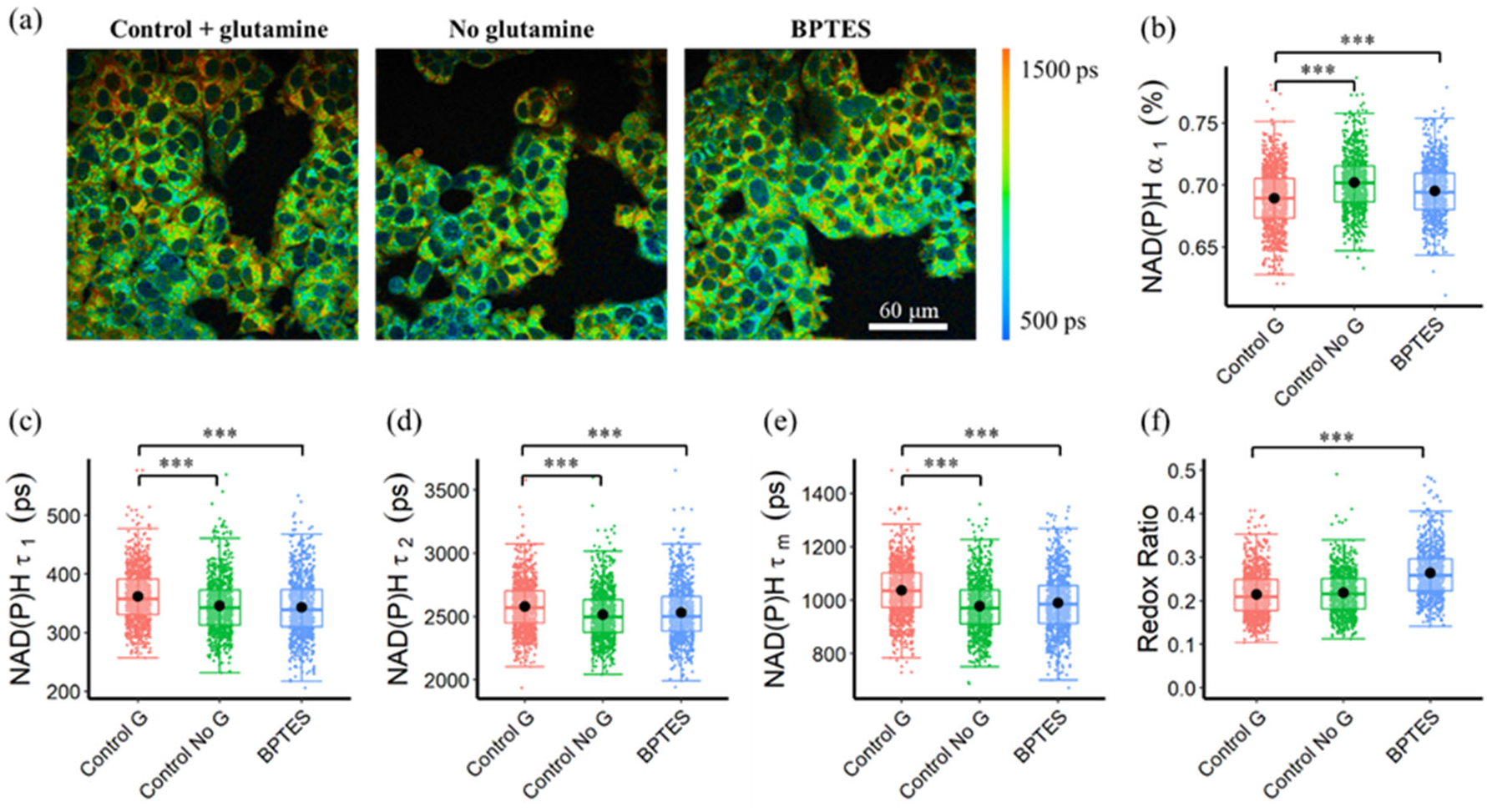
Autofluorescence lifetime variations of MCF7 cells in response to glutaminolysis inhibition. (a) Representative NAD(P)H *τ_m_* images of control (Control G), no glutamine (Control No G), and BPTES-treated (BPTES) MCF7 cells, scale bar = 60 μm (b) NAD(P)H *α_1_* (c) NAD(P)H *τ_1_* (d) NAD(P)H *τ_2_* (e) NAD(P)H *τ_m_* (f) Intensity redox ratio (FAD/ (FAD + NAD(P)H)) of MCF7 cells in response to glutaminolysis perturbations. ***P < 0.001 for two-sided student t test with Bonferroni correction for multiple comparisons. Substrates in each media: Control G (25 mM glucose + 1 mM pyruvate + 2 mM glutamine), Control No G (25 mM glucose + 1 mM pyruvate), BPTES (25 mM glucose + 1 mM pyruvate+ 2 mM glutamine + 10 μm BPTES)

To isolate the effects of glutaminolysis from OXPHOS which proceeds once a cell has converted glutamine to *α*-ketoglutaric acid, autofluorescence lifetime imaging was performed on MCF7 cells fasted for 1 hour and then exposed to 2 mM glutamine. An increase in NAD(P)H intensity and FAD intensity within the MCF7 cells was observed at 2 and 3 hours of glutamate, as compared with the cells with glutamate for 1 hour (Sup Fig 5). The NAD(P)H and FAD fluorescence lifetime components changed over time with glutamate stimulus. 1 hour of glutamine stimulus increased the NAD(P)H lifetimes (*τ_1_*, *τ_2_*, *τ_m_*) and reduced the free fraction (*α_1_*) of NAD(P)H as compared with control MCF7 cells (Sup Fig 6(a)(b)(c)(d)). Similarly, as compared with control MCF7 cells, 1, 2, and 3 hours of glutamine stimulus increased the FAD lifetimes (*τ_1_*, *τ_2_*, *τ_m_*) and reduced the bound fraction (*α_1_*) of FAD (Sup Fig 6(e)(f)(g)(h)).

### Pyruvate concentration alters the FAD fluorescence lifetime

To evaluate the effects of OXPHOS stimulation on autofluorescence lifetime metrics, MCF7 cells were fasted and then provided pyruvate at scaled concentrations. Cellular quantitation analysis showed that the pyruvate concentration groups had a longer bound (*τ_1_*) and free (*τ_2_*) FAD lifetime, which led to a longer mean FAD lifetime (*τ_m_*), as compared to the control group (Figure 4(c)(d)(e)). The cells exposed to different concentrations of pyruvate had an increased fraction of enzyme-bound FAD (*α_1_*) than the control cells (Figure 4(b)). Furthermore, pyruvate starvation caused a reduced fraction of bound FAD (*α_1_*), while increased pyruvate concentration increased the bound FAD fraction (*α_1_*), and bound FAD lifetime (*τ_1_*) (Figure 4(b)(c)). A longer mean NAD(P)H fluorescence lifetime (*τ_m_*) of MCF7 cells can be observed in the pyruvate groups than in the control groups due to reduced free NAD(P)H fraction, and increased free and bound NAD(P)H lifetime (Sup. Fig 7(c)(d)(e)(f)). Additionally, the intensity of both NAD(P)H and FAD were lower in the pyruvate assay groups than in the control group, and an increased pyruvate concentration reduced both the NAD(P)H and FAD intensities, which led to reduced optical redox ratio (Figure 4(f)(g), Sup Fig 7(b)).

**Figure 4.**
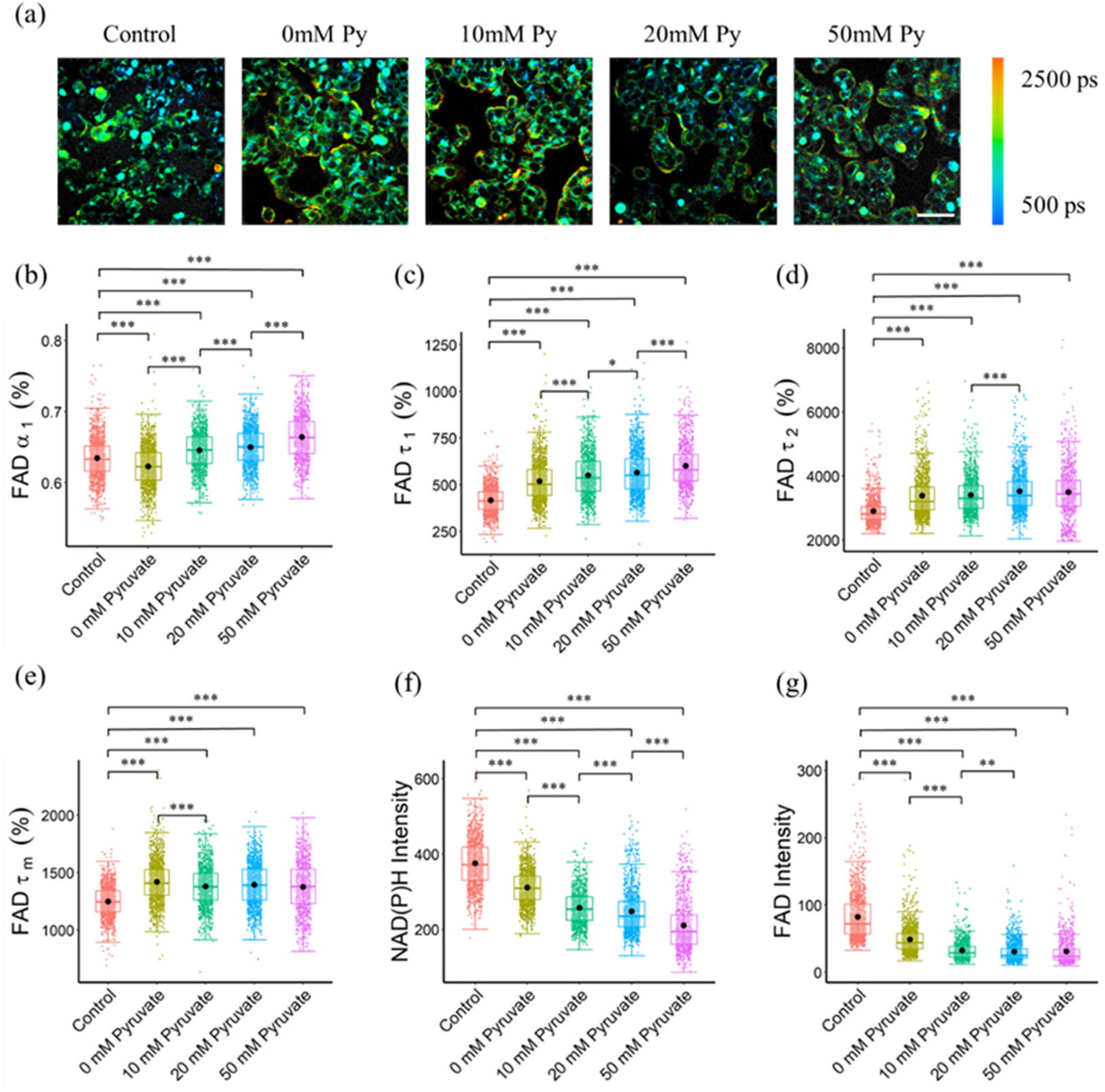
FAD fluorescence lifetimes of MCF7 cells were altered with different pyruvate concentrations of the culture media. (a) Representative FAD *τ_m_* images of cancer cells exposed to different pyruvate concentrations, Py, pyruvate; scale bar = 60 *μm* (b) FAD *α_1_* (c) FAD *τ_1_* (d) FAD *τ_2_* (e) FAD *τ_m_* (f) NAD(P)H intensity and (g) FAD intensity of MCF7 cells with different pyruvate concentrations. *P < 0.05, **P < 0.01, ***P < 0.001 for two-sided student t test with Bonferroni correction for multiple comparisons. Substrates in each media: Control (25 mM glucose + 1 mM pyruvate), Pyruvate (no glucose + no glutamine + 0/10/20/50 mM pyruvate).

### NAD(P)H and FAD fluorescence lifetime features predict cellular metabolic perturbations

UMAP visualization of the 12 autofluorescence imaging features (NAD(P)H *α_1_*, *τ_1_*, *τ_2_*, *τ_m_*, intensity; FAD *α_1_*, *τ_1_*, *τ_2_*, *τ_m_*, intensity; redox ratio; FLIRR) revealed the cluster separation of MCF7 cells with glycolysis inhibition groups (grey dots) from the OXPHOS inhibition groups (blue dots) with the control cell group (red dots) in the middle and overlapping with both inhibition groups (Figure 5(a)). A UMAP of the subset of data with MCF7 cells treated with 50 mM 2-DG and glucose starvation as the glycolysis inhibition groups, and treated with cyanide for maximizing the OXPHOS activities, showed a significant separation of MCF7 cells with inhibited glycolysis (grey dots) and cells with inhibited OXPHOS (blue points) (Figure 5(b)).

**Figure 5.**
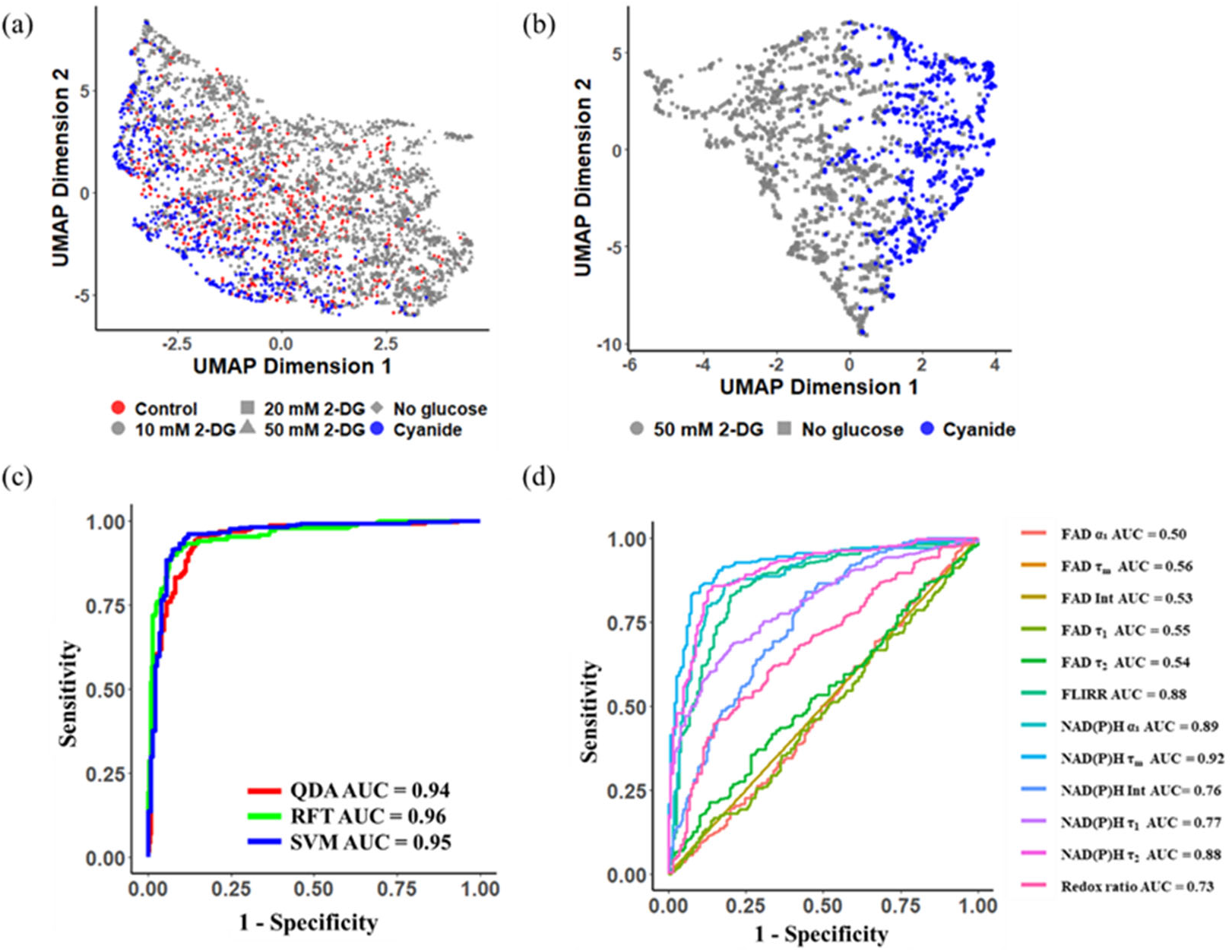
Autofluorescence lifetime features allowed classification of metabolic perturbations of MCF7 cells (a) (b) UMAP data reduction technique allows visual representation of the separation between OXPHOS maximum (glycolysis inhibition groups: 10 mM 2-DG, 20 mM 2-DG, 50 mM 2-DG, No glucose) and glycolysis maximum cells (OXPHOS inhibition groups: Cyanide). Each color represents a metabolic group, red corresponds to control, grey and blue to glycolysis inhibition, and OXPHOS inhibition respectively. Each shape represents a different drug-treated group. ROC curves of the test data for (c) machine learning classification models, RFT: random forest tree; SVM: support vector machine; QDA: quadratic discriminant analysis, and (d) lifetime features for classification of glycolytic versus oxidative cancer cells. Int, intensity; Redox ratio=FAD/(FAD + NAD(P)H); FLIRR=NAD(P)H *α_2_* FAD *α_1_*

Using the MCF7 metabolic perturbation data, machine learning algorithms were developed to predict cellular metabolic variations. A random forest tree (RFT) model achieved a mean prediction accuracy of 92% and an ROC AUC value of 0.96 (Figure 5(c), Sup Fig 8(a)). The support vector machine (SVM) model achieved a mean prediction accuracy of 90% and an ROC AUC of 0.95 (Figure 5(c), Sup Fig 8(b)). The quadratic discriminant analysis (QDA) obtained a mean prediction accuracy of 92% and an AUC value of 0.94 (Figure 5(c), Sup Fig 8(c)). When normalizing the data with the corresponding control cell group, the random forest tree model achieved a mean prediction accuracy of ~ 0.97 with an ROC AUC value of ~ 0.99 (Sup Fig 8(d)(e)).

Feature analysis from the random forest tree model revealed that the mean NAD(P)H lifetime (*τ_m_*) contributed most to this prediction, followed by the free NAD(P)H fraction (*α_1_*) (Sup Fig 9). Furthermore, the ROC AUC of models built from each feature individually implied that NAD(P)H *τ_m_* (AUC = 0.92), and NAD(P)H *α_1_* (AUC = 0.89), FLIRR (AUC = 0.88) and NAD(P)H *τ_2_* (AUC = 0.88) were significant features for the prediction of cancer cell metabolic perturbations (Figure 5(d)). FAD lifetime features had low ROC AUC values and feature importance (Figure 5(d), Sup Fig 9).

### Convolutional neural networks can predict metabolic activities of MCF7 cells from NAD(P)H and FAD fluorescence lifetime images

We trained a conventional LeNet CNN architecture with different autofluorescence lifetime feature images and tuned the training parameters to achieve the best performance. The performance of each CNN model was assessed based on the prediction results of the test dataset. The CNN models reached the least validation loss at ~ 80 epochs with a 0.00001 learning rate (Sup. Fig 10). When trained with only NAD(P)H intensity images, the LeNet CNN achieved an average test dataset accuracy of 75.7%, an ROC AUC of 0.81, a precision of 76.5%, and a recall of 60.0% for predicting glycolytic versus oxidative cells (Table 3, Figure 6(a)(b)). Similar to the classical machine learning model results, including FAD intensity images did not significantly improve this prediction. When the CNN model was trained on both NAD(P)H intensity and FAD intensity images together, the accuracy of the test data increased by ~3% (78.4%) with an ROC AUC value of 0.86 (Table 3, Figure 6(b)). The CNN model performed better when trained with NAD(P)H lifetime images, as compared to the intensity images. The CNN model trained with NAD(P)H *τ_m_* images classified glycolytic MCF7 cells from oxidative cells with an average accuracy of 85.1%, a precision of 87.4%, a recall of 81.3%, and an ROC AUC of 0.93 for the test data (Table 3, Figure 6(a)(b)). Including additional NAD(P)H lifetime component images in the CNN improved the accuracy, and the highest performance achieved for the CNN model was trained with all NAD(P)H lifetime components (intensity, *τ_1_*, *τ_2_*, *τ_m_*, *α_1_*). This model achieved 94.5% accuracy, 0.99 ROC AUC, 98.2% precision, and 91.5% recall. (Table 3, Figure 6(a)(b)). The performance of the LeNet CNN trained with images of all NAD(P)H lifetime components (intensity, *τ_1_*, *τ_2_*, *τ_m_*, *α_1_*) exceeded the prediction performance of classical machine learning algorithms, including RFT, SVM, and QDA, trained and tested with the same datasets (Table 3).

**Table 3.** Performance metrics for the test dataset (n = 1520 cells) of each classifier built to predict MCF7 cell metabolism as glycolysis inhibition or OXPHOS inhibition. LeNet CNN models were trained on cropped images of individual cells. RFT, SVM, and QDA models were trained on the lifetime features of the cells averaged across the cellular pixels.

| Model | Accuracy | AUC | Precision | Recall |
| --- | --- | --- | --- | --- |
| CNN (NAD(P)H Intensity) | 75.7% | 0.81 | 76.5% | 60.0% |
| CNN (NAD(P)H $\tau_m$) | 85.1% | 0.93 | 87.4% | 81.3% |
| CNN (NAD(P)H Intensity + FAD Intensity) | 78.4% | 0.86 | 82.6% | 60.6% |
| CNN (NAD(P)H Intensity + $\tau_m$) | 89.9% | 0.96 | 95.2% | 84.5% |
| CNN (NAD(P)H $\alpha_1$ + $\tau_1$ + $\tau_2$ + $\tau_m$ + Intensity) | 94.5% | 0.99 | 98.2% | 91.5% |
| RFT (NAD(P)H $\alpha_1$ + $\tau_1$ + $\tau_2$ + $\tau_m$ + Intensity) | 88.6% | 0.95 | 92.4% | 87.3% |
| SVM (NAD(P)H $\alpha_1$ + $\tau_1$ + $\tau_2$ + $\tau_m$ + Intensity) | 86.6% | 0.93 | 89.0% | 85.1% |
| QDA (NAD(P)H $\alpha_1$ + $\tau_1$ + $\tau_2$ + $\tau_m$ + Intensity) | 87.1% | 0.94 | 90.4% | 88.7% |

RFT: random forest tree; SVM: support vector machine; QDA: quadratic discriminant analysis.

**Figure 6.**
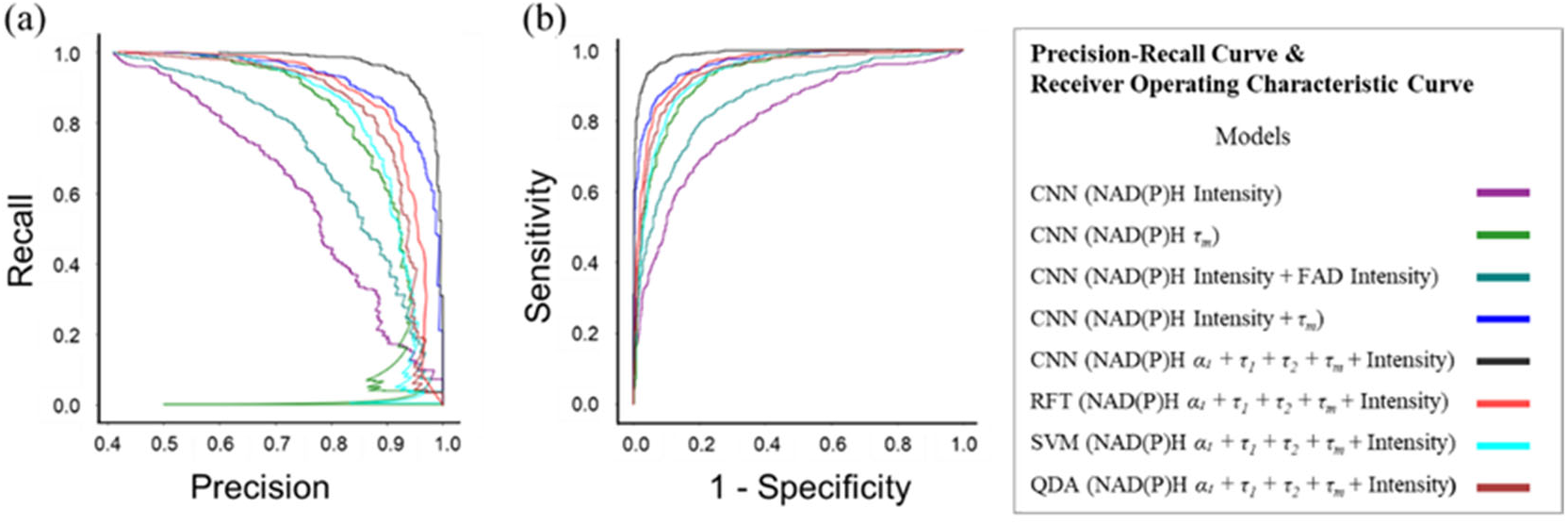
Prediction performance of classifiers to predict MCF7 metabolism as glycolysis inhibition or OXPHOS inhibition from autofluorescence lifetime images. (a) Precision-recall curves of each classifier. (b) AUC ROC curves for each classifier. RFT: random forest tree; SVM: support vector machine; QDA: quadratic discriminant analysis

### The metabolic prediction models transfer to additional datasets

To further access the applicability of the fluorescence lifetime model to predict metabolic pathways, we firstly tested the models with data from the 10 mM and 20 mM 2-DG concentration groups, and the pyruvate titration data. Using the classical machine learning model, more than 80% of the MCF7 cells exposed to 10 mM 2-DG and 20 mM 2-DG were predicted to have glycolysis inhibition (Table 4). When tested with the pyruvate assay data, where MCF7 cells were only provided pyruvate as a substrate, more than 95% of the cells were predicted to have glycolysis inhibition by the classical machine learning model (Table 4).

**Table 4.** Prediction results from the normalized random forest tree model for MCF7 cell metabolism for the 10 mM 2-DG, 20 mM 2-DG, 0 mM pyruvate, 10 mM pyruvate, 20 mM pyruvate, and 50 mM pyruvate groups. 2-DG inhibits glycolysis. Pyruvate stimulates OXPHOS.

|  | 10 mM 2-DG | 20 mM 2-DG | 10 mM Pyruvate | 20 mM Pyruvate | 50 mM Pyruvate |
| --- | --- | --- | --- | --- | --- |
| Glycolysis Inhibition | 82.6% (659) | 85.9% (531) | 98.5% (1152) | 98.9% (1217) | 99.2% (924) |
| OXPHOS Inhibition | 17.4% (139) | 14.1% (87) | 1.5% (17) | 1.1% (13) | 0.8% (7) |

Finally, we evaluated if the models trained with MCF7 cells can be used to identify metabolic variations of other cells. Glucose-treated HepG2 liver cancer cells had more fraction of free NAD(P)H (*α_1_*), and shorter NAD(P)H lifetimes (*τ_m_*) than the control cells (Figure 7(a), Sup. Fig 11(a)(d)). Palmitate treatment of HepG2 cells caused a lower fraction of free NAD(P)H (*α_1_*), and longer free and bound NAD(P)H lifetimes (*τ_1_*, *τ_2_*) compared to the control cells (Sup. Fig 11(a)(b)(c)). When adapting the UMAP algorithm to visualize distributions of different cellular groups, a separation between cells exposed to glucose and cells exposed to palmitate (PA) was observed (Figure 7(b)). When applying the RFT machine learning model trained with normalized MCF7 cell data, 65.4% of the glucose-treated cells were predicted to be glycolytic (OXPHOS inhibition), and 89.2% of the palmitate exposed cells were predicted to be oxidative (glycolysis inhibition) (Figure 7(c)). When applying the CNN model (trained with all NAD(P)H lifetime features) that was trained with the MCF7 cells to the HepG2 liver cancer data, 80.8% of the palmitate exposed cells were predicted to have glycolysis inhibition which was consistent with the prediction of the classical machine learning model. In contrast, 64.4% of the glucose-exposed cells were predicted to be glycolysis inhibition (Figure 7(d)). Moreover, MCF7 metabolic prediction models were also tested with FLIM data from primary human T cells, which have different metabolic phenotypes by activation state. Activated T cells are more dependent on glycolysis and quiescent T cells are more dependent on OXPHOS. Using the random forest tree model, 80.2% of the activated T cells were predicted to be glycolytic (OXPHOS inhibition), and 97.6% of the quiescent T cells were predicted to be oxidative (glycolysis inhibition) (Figure 7(e)). However, most (98+%) of both the activated and quiescent T cells were predicted to exhibit OXPHOS inhibition with the CNN model (Sup. Table 4).

**Figure 7.**
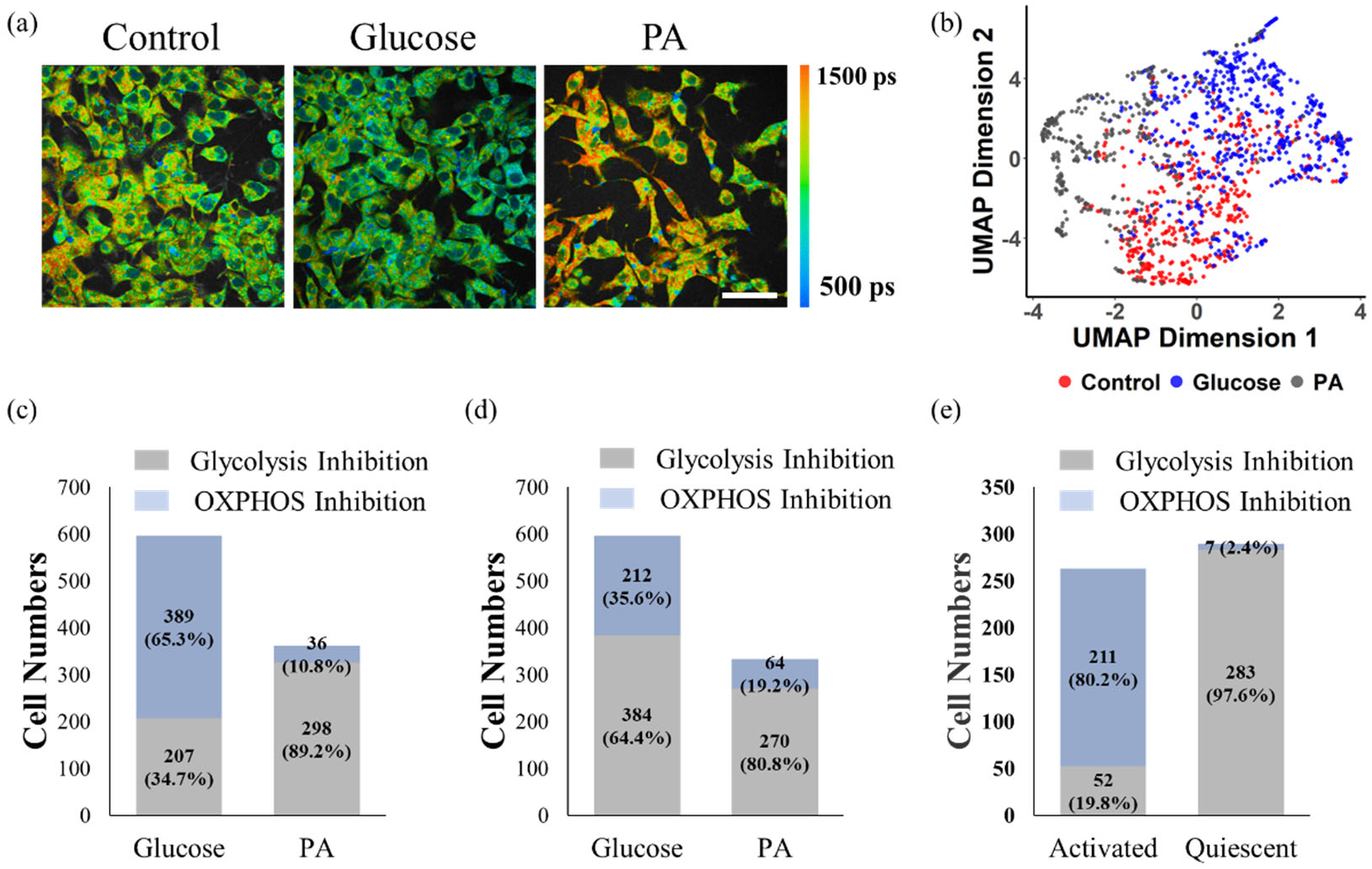
MCF7 cell-trained model prediction performance for liver cancer cells and T cells. (a) Representative NAD(P)H *τ_m_* images of HepG2 cells exposed to control media (starved condition), glucose (30 mM), and palmitate (0.4 mM PA), scale bar = 60 μm (b) UMAP data reduction technique allows visual representation of the separation between different metabolic groups of hepatocellular cells. Each color represents a metabolic group, red corresponds to control, grey and blue to glucose treated, and PA treated respectively. (c) Prediction of HepG2 cell metabolism by the random forest tree model trained with MCF7 cells. (d) Prediction of HepG2 cell metabolism by the CNN model trained with MCF7 cells. (e) Prediction of activated and quiescent T cell metabolism by the random forest tree model trained with MCF7 cells.

## Discussion

Autofluorescence lifetime imaging is sensitive to metabolic differences in live cells between groups such as cancer and non-cancer cells, phenotypes of immune cells, and stem cells and differentiated cells (20, 25–29, 32–35). By using endogenous fluorophores for contrast, autofluorescence lifetime imaging resolves cellular and sub-cellular resolution without contact or manipulation of the sample providing label-free advantages and independence from label-related confounding factors. However, autofluorescence measurements lack the specificity provided by protein- or molecule-targeted labels. Selection of specific excitation and emission wavelengths can isolate endogenous fluorophores such as NAD(P)H and FAD (37). Two-component exponential decay fitting of NAD(P)H and FAD fluorescence lifetimes allows quantification of the fraction of bound or free coenzymes, and the lifetime values of the short and long lifetimes (23, 38). Despite this biochemical specificity, an interpretation or relationship between NAD(P)H and FAD fluorescence lifetime metrics and macroscopic cellular metabolic phenotypes remains elusive. In this study, we designed metabolic perturbation experiments to selectively activate or inhibit the metabolic pathways of glycolysis, OXPHOS, and glutaminolysis via controlled substrate availability and metabolic inhibitors. Then, NAD(P)H and FAD fluorescence lifetime images were acquired and analyzed to define fluorescence lifetime features of cells using specific metabolic pathways. Finally, this robust dataset of autofluorescence lifetime data was used to design and evaluate models for predicting cellular use of glycolysis or OXPHOS. The results show that fluorescence lifetime imaging of NAD(P)H and FAD allows a non-destructive technique to predict metabolic perturbations of cancer cells when treated with inhibitors or exposed to culture environment variations at the single-cell level.

Changes in metabolic pathways within MCF7 cells alter the fluorescence lifetimes of NAD(P)H and FAD. Cancer cells often have enhanced glycolysis, even in the presence of oxygen and uncompromised mitochondrial function, to promote growth, proliferation, and survival (1). To isolate the effects of glycolysis on autofluorescence lifetimes, MCF7 cells were exposed to cyanide, which inhibits oxidative phosphorylation as validated by the reduction of OCR in the seahorse experiment (Figure 8, Sup Fig 3(b)). Glycolysis, in the absence of OXPHOS due to cyanide, results in an increased fraction of free NAD(P)H and reduced free and bound NAD(P)H lifetimes (Figure 2, Table 2). Conversely, glycolysis inhibition with 2-DG treatment and OXPHOS stimulation with glucose starvation and pyruvate stimulation result in the exact opposite changes in NAD(P)H lifetime features: a reduced fraction of free NAD(P)H and increases of the NAD(P)H lifetimes (Figure 2, Table 2, Sup Fig 7). The reduction of free NAD(P)H fraction (*α_1_*) with 2-DG treatment has also been observed in MCF10A breast cancer cells and pancreatic islet cells (39, 40). Conversely, a higher level of glycolysis induces more fraction of free NAD(P)H in kidney cells, neural cells, and stem cells in disease states (41–43). The consistent and opposite changes in NAD(P)H fluorescence lifetimes support the hypothesis that NAD(P)H fluorescence lifetimes are responsive to metabolic pathway shifts and changes in the protein binding partners of NAD(P)H (44, 45). Consequently, NAD(P)H *τ_m_* was identified as the most important feature for the differentiation of glycolytic and oxidative cells across various feature evaluation methods (Sup Fig. 9, Figure 5(d)). Furthermore, NAD(P)H *α_1_* also contributes significantly to the prediction of glycolytic versus oxidative cells (Figure 5(d), Sup Fig. 9). NAD(P)H *α_1_* is the highest weighted feature for the classification of T cell activation, allowing robust differentiation of oxidative quiescent T cells from glycolytic, activated T cells (35).

**Figure 8.**
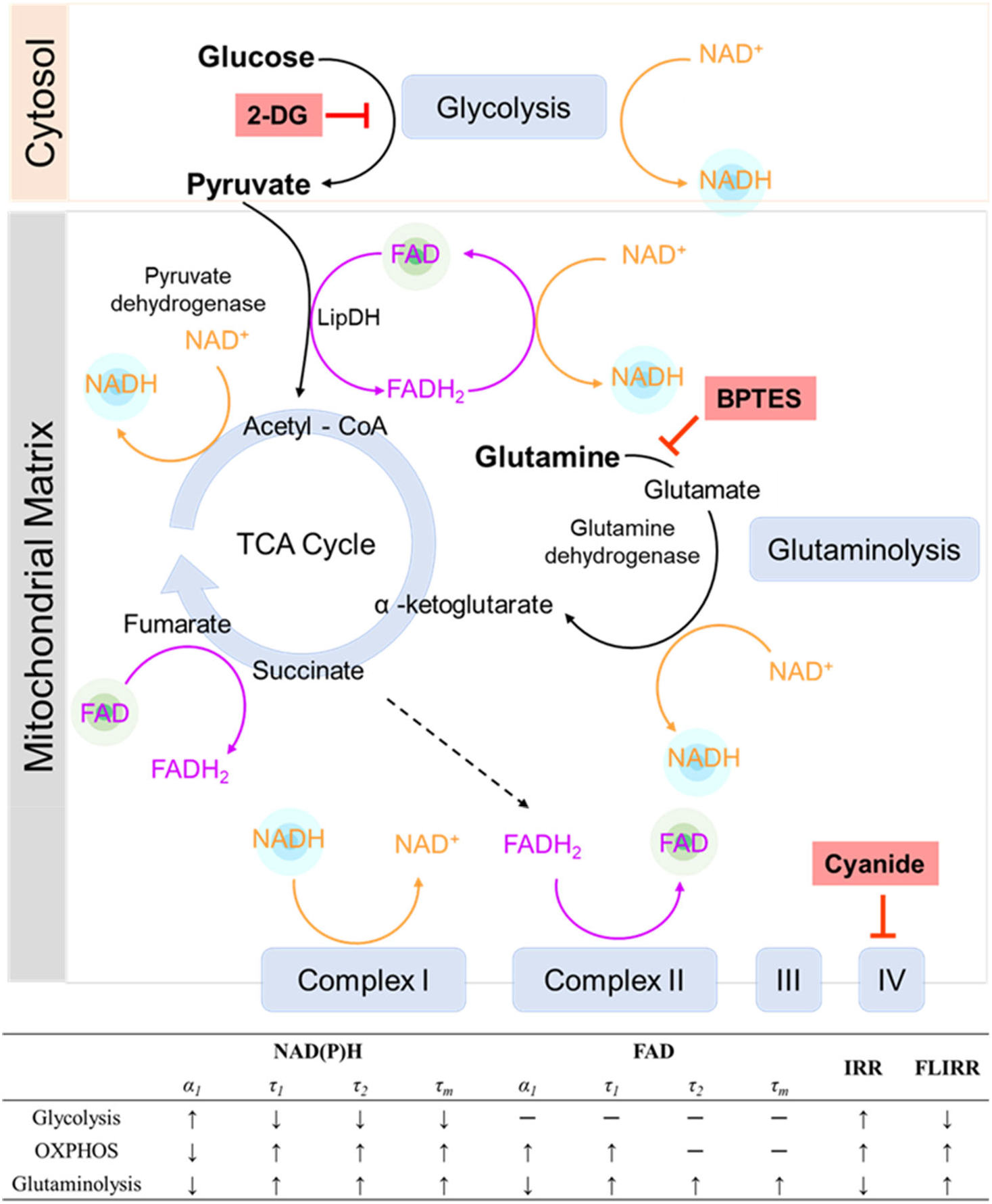
The roles of NADH and FAD in metabolic pathways. NADH and FAD are coenzymes in glycolysis, the TCA cycle, glutaminolysis, and the electron transport chain. 2-DG inhibits glycolysis, forcing a cell to use alternative pathways for metabolism. Cyanide inhibits complex IV of the electron transport chain, effectively inhibiting OXPHOS. BPTES inhibits glutaminolysis. Each metabolic pathway varies the fluorescence lifetimes of NAD(P)H and FAD differently. IRR = intensity redox ratio FAD/(FAD+NAD(P)H), FLIRR = fluorescence lifetime redox ratio NAD(P)H *α_2_*/ FAD *α_1_*

In contrast to the consistent and opposite variations observed in NAD(P)H lifetime features of MCF7 cells with isolated glycolysis and OXPHOS metabolism (Figure 8), perturbations of glycolysis and OXPHOS with 2-DG, glucose starvation, and cyanide did not significantly and consistently alter FAD fluorescence lifetime metrics of MCF7 cells (Table 2, Sup Fig. 2). Consequently, the FAD lifetime features contribute less than the NAD(P)H lifetime features to the metabolic pathway prediction model (Sup Fig. 9, Figure 5(d)). These findings are inconsistent with published FAD lifetime changes in lung cancer cells exposed to rotenone/antimycin for OXPHOS inhibition, which results in a reduced free-to-bound fraction of FAD (46). Additionally, multiple variations of FAD fluorescence lifetime have been observed in bladder cancer cells, skin cells, stem cells, and neural cells due to shifts between glycolysis and OXPHOS (42, 47–49). The use of FAD as a primary electron carrier at multiple steps throughout the TCA and the electron transport chain suggests that FAD fluorescence lifetimes should be sensitive to alterations in cellular use of the oxidative metabolic pathway. To further evaluate the role of OXPHOS on FAD fluorescence lifetimes, MCF7 cells were grown and imaged in media with titrated concentrations of pyruvate and an absence of other metabolic substrates. Pyruvate enters the mitochondria and is converted to acetyl-CoA by pyruvate dehydrogenase (PDH) to facilitate oxidative phosphorylation (Figure 8). In this process, pyruvate dehydrogenase (PDH) acts as a gatekeeper between glycolysis and oxidative phosphorylation, and FAD is reduced to FADH_2_ through lipoamide dehydrogenase (LipDH) (Figure 8). FADH_2_ can then be oxidized to reduce NAD^+^ to NAD(P)H, which contributes to electron transport chain. While a 1 mM stimulus of pyruvate did not alter the OCR of MCF7 cells, the NAD(P)H intensity and optical redox ratio ((FAD/(NAD(P)H + FAD)) were reduced at 10, 20, and 50 mM pyruvate (Sup Fig 1, Figure 4(f), Sup Fig 7), which implies an increased oxidative state within the cells (29). An increased fraction of bound FAD of MCF7 cells is observed with increased pyruvate media concentrations than in the control cells (Figure 4(b)), consistent with the expected increased use of FAD due to increased oxidative metabolism and throughput in the TCA cycle, where FAD catalyzes the oxidation of succinate into fumarate (Figure 8) (50).

Elevated levels of glutaminolysis have also been found in cancer cells to compensate for changes in glycolysis and maintain a functional TCA cycle (51–53). Glutamine is converted to glutamate by glutaminase, and further metabolized to *α*-ketoglutarate by glutamate dehydrogenase and fuels the TCA cycle (Figure 8). In glutaminolysis, NAD^+^ is reduced to NADH and contributes to the mitochondrial-bound NADH pools (29) (Figure 8). To isolate the effects of glutaminolysis on the fluorescence lifetimes of NAD(P)H and FAD, the glutaminolysis pathway of MCF7 cells was stimulated by fasting and then stimulating with glutamine (2 mM) and inhibited by exposure to BPTES. Fluorescence lifetimes of NAD(P)H and FAD were tracked over time following glutamine stimulation and an increased NAD(P)H fluorescence intensity was observed at 1, 2, and 3 hours (Sup Fig 5(a)), consistent with the expectation that glutaminolysis increases NADH. The contribution of glutaminolysis to the bound NAD(P)H fraction in MCF7 cells are consistent with prior published results in human foreskin keratinocytes and C2C12 myoblasts (54). In an absence of glutaminolysis, it is expected that NADH would not be generated through this pathway and lead to an increased optical redox ratio (FAD/(NAD(P)H+FAD), which was observed in the BPTES-treated MCF7 cells (Figure 3(f)). Furthermore, inhibition of glutaminolysis by BPTES has also been found to decrease ATP levels and profoundly increase ROS levels (55). Glycolysis and glutaminolysis elicit opposite changes in structural metabolic readouts, including mitochondrial clustering and network analysis (54, 56, 57). Similarly, glutaminolysis induces opposite trends of NAD(P)H fluorescence lifetime metrics of MCF7 cells as glycolysis (Figure 8).

Fluorescence lifetime imaging is independent of experimental factors including laser power, filter effects, and detector sensitivity, and provides a more robust result than fluorescence intensity imaging. Additionally, fluorescence lifetime imaging also provides information regarding binding status and microenvironment of the fluorophores, which can be used to infer additional biochemical information about NAD(P)H and FAD. Therefore, the FLIRR, a metric of bound NAD(P)H to bound FAD, was created to overcome the limitations of the traditional intensity-based redox ratio (IRR), yet provide metabolic information within a single metric (46, 58). The FLIRR is sensitive to drug-induced metabolic alterations of human keratinocytes, and squamous cancer cells (59). FLIRR reflects the protein-bound ratio of NAD(P)H and FAD and is not correlated with IRR across a variety of metabolic states in T cells (60). The intensity redox ratio (IRR) can be calculated in different formats with FAD and NAD(P)H as the numerator and normalization, and these computations are equivalent in detecting T cell metabolic perturbations (60). Additionally, the FAD numerator redox ratio (FAD/ (FAD + NAD(P)H)) is sensitive to dynamic changes in oxygen consumption rate of breast cancer cells (61). Our results show that both glycolysis and OXPHOS metabolic activities lead to a higher IRR (FAD/(FAD + NAD(P)H)), but glutaminolysis reduces the IRR (Figure 8). In contrast, glycolysis induces a lower FLIRR, but OXPHOS and glutaminolysis stimulate a higher FLIRR (Figure 8).

Due to the unique autofluorescence lifetime phenotypes observed for glycolytic or oxidative MCF7 cells, we compared different models for predicting cellular metabolic phenotypes from the NAD(P)H and FAD fluorescence lifetime imaging data. Machine learning models are appropriate for analysis of datasets with multiple variables, and have been used to extract and interpret cell phenotypes from fluorescence lifetime data. Several studies have applied extracted lifetime features with machine learning algorithms to identify mouse embryo health, quantify precancer cells, classify T cell activation, differentiate stem cell phenotypes, and investigate metabolic perturbations (35, 54, 62–66). Here, both conventional machine learning methods and neural networks were trained with autofluorescence lifetime images and features to predict metabolic states of cancer cells. High ROC AUCs (~0.95) and accuracy (~0.91) were achieved for test datasets evaluated by machine learning algorithms using autofluorescence imaging features (intensity redox ratio, FLIRR, NAD(P)H *τ_m_*, NAD(P)H *τ_1_*, NAD(P)H *τ_2_*, NAD(P)H *α_1_*, NAD(P)H intensity, FAD *τ_m_*, FAD *τ_1_*, FAD *τ_2_*, FAD *α_1_*, and FAD intensity) to classify MCF7 cells as using OXPHOS or glycolysis (Figure 5(c)). The accuracy of the model is comparable with published papers that use NAD(P)H intensity images or autofluorescence lifetime features to predict cell phenotypes (45, 65, 66). Normalization with the mean lifetime values of the control group improves classification accuracy (0.97), suggesting heterogeneity within the cancer cells or across technical replicates can be reduced by normalization (Sup Fig 8).

The mitochondrial network structure is associated with metabolism and cellular function (67–71). Since NAD(P)H is primarily localized in mitochondria and thus mitochondria appear as bright pixels in NAD(P)H images, analyzing mitochondrial organization from autofluorescent images offers complimentary morphology information, and reflects alterations in metabolic activities (72, 73). Specifically, the mitochondria are observed to be more clustered with a higher glycolytic level and less clustered when glutaminolysis and OXPHOS are dominant in tissues and cells (45, 56). The high resolution of autofluorescence lifetime imaging provides subcellular information and allows insights for the measurement at the single-cell level. However, obtaining the autofluorescence lifetime features by averaging the pixel value in the cytoplasm regions disregards the subcellular information related to the metabolic status. The CNN can extract hidden patterns in the images automatically, but requires a larger dataset for training as compared to conventional machine learning methods. The LeNet CNN model achieved around 89% accuracy in classifying T cell activation when trained with NAD(P)H intensity images (66). However, the information from the NAD(P)H intensity images alone classifies glycolysis versus OXPHOS in MCF7 cells with only 79.5% accuracy (Table 3). Including all NAD(P)H fluorescence lifetime component images together improves the accuracy of the LeNet CNN to 95% accuracy and an AUC of 0.99 for the testing dataset (Table 3, Figure 6). The CNN model can automatically learn features from the images and exceeds the performance of conventional machine learning models that use a single extracted FLIM feature for each cell, confirming the added value of retaining spatial information (Table 3). However, the recall of the CNN model is lower than its precision, suggesting higher false-negative predictions which might be due to heterogeneity within the cells (Table 3, Figure 6(a)).

To ensure the utility of the metabolism-prediction models for additional cells and studies beyond MCF7 cells, we tested the models trained with breast cancer cells on different samples. For liver cancer cells, palmitate treatment triggers fatty acid oxidation, which produces acetyl-CoA and contributes to the TCA cycle and electron transport chain. Palmitate has been observed to reduce the NAD(P)H bound fraction in myoblast cells, which is in contrast to the cancer cells (45). Even though there are multiple metabolic pathways besides OXPHOS happening in the cancer cells with PA treatment, our model only predicts whether glycolysis or oxidation is the major metabolic activity. The pre-trained CNN model did not provide a good result when used to determine the metabolic phenotypes of liver cancer cells and T cells (Figure 7(d), Sup. Table. 3), implying that the difference in morphological features such as cell size, and subcellular structure influence the performance of the CNN significantly. In contrast, the conventional machine learning models that use averaged FLIM features for each cell rather than the cell images, worked well to predict glycolysis or OXPHOS use by hepatocellular cells, suggesting that the autofluorescence endpoints reflect changes in metabolic variations that are consistent across these two different cancer cell types (Figure 7(c)). Furthermore, the T cell prediction results of the RFT model trained on MCF7 data (Figure 7(e)) revealed that the metabolic-prediction model was also useful for identifying the metabolic switch of T cells switch from oxidative phosphorylation to glycolysis and glutaminolysis upon activation (74, 75). Specifically, the T cell data was collected on a different microscope at a different location, indicating that the metabolic-prediction model can be used across cell types, phenotypes, and different lifetime microscopes. The applicability of the MCF7-cell trained RFT model to accurately predict metabolic phenotypes across other cells and metabolic perturbations supports the promotion of ML-FLIM for identifying cellular metabolism in extensive research fields.

In this paper, unique NAD(P) and FAD lifetime alterations were observed within cancer cells with different levels of glycolysis, OXPHOS, and glutaminolysis. These alterations are sufficient for machine learning models to predict the predominance of glycolysis versus OXPHOS from the lifetime features and images. The FLIM image-based CNN models provided increased accuracy over traditional feature-based machine learning models due to spatial information within the cell, but are less transferrable to other cells. The RFT model trained with MCF7 cells can accurately predict the dominant metabolic pathway of liver cancer cells exposed to different metabolic environments and the metabolic status of activated or quiescent T cells. In summary, autofluorescence lifetime imaging offers a label-free, quantitative method to identify metabolic activities in living cells, and can be used across various platforms for broad applications to study metabolic changes due to chemotherapy, gene expression, and immune responses.

## Acknowledgments

Texas A&M University School of Medicine Analytical Cytometry Core for the use of the Seahorse Analyzer. Funding sources include NIH NIGMS R35 GM142990 (AJW).

## Author Contributions

The hypothesis and methodology were conceptualized by AJW and LH. Cell imaging experiments were performed by LH, NW, and LL, with supervision by AJW and LX. All data analysis was performed by LH, and machine learning models were developed by LH with supervision by AJW. The seahorse metabolic flux assay experiment was performed by LH, NW, and JB, with supervision by APW. A preliminary manuscript draft was written by LH. All authors contributed to the reviewing and editing of the manuscript.

