## Supplementary Materials for "Machine learning prediction of cancer cell metabolism from autofluorescence lifetime images"

Supplementary Table 1 The concentration of metabolic substrates in each group

| Media | The concentration of metabolic substrates |
|---|---|
| Control Media 1 (No glutamine) | glucose (25 mM), pyruvate (1 mM), no glutamine |
| 10 mM 2-DG | glucose (25 mM), pyruvate (1 mM), no glutamine, 2-DG (10 mM, 1 hour) |
| 20 mM 2-DG | glucose (25 mM), pyruvate (1 mM), no glutamine, 2-DG (20 mM, 1 hour) |
| 50 mM 2-DG | glucose (25mM), pyruvate (1 mM), no glutamine, 2-DG (50 mM, 1 hour) |
| No glucose | no glucose, pyruvate (50 mM, 1 hour), no glutamine |
| Cyanide | glucose (25mM), pyruvate (1 mM), no glutamine, NaCN (4 mM) |
| Control Media 2 (0 mM pyruvate) | no glucose, no pyruvate, no glutamine |
| 10 mM pyruvate | no glucose, pyruvate (10 mM, 1 hour), no glutamine |
| 20 mM pyruvate | no glucose, pyruvate (20 mM, 1 hour), no glutamine |
| 50 mM pyruvate | no glucose, pyruvate (50 mM, 1 hour), no glutamine |
| Control Media 3 | glucose (25mM), pyruvate (1 mM), glutamine (2 mM) |
| BPTES | glucose (25 mM), pyruvate (1 mM), glutamine (2 mM), BPTES (10 µm, 1 hour) |
| Glutamine 1 | no glucose, no pyruvate, glutamine (2 mM, 1 hour) |
| Glutamine 2 | no glucose, no pyruvate, glutamine (2 mM, 2 hours) |
| Glutamine 3 | no glucose, no pyruvate, glutamine (2 mM, 3 hours) |

Supplementary Table 2 Number of liver cancer cells and T cells

| Liver cancer cells | Cell Number | T cells | Cell Number |
|---|---|---|---|
| Control | 451 | Activated T cells | 263 |
| Glucose | 596 | Quiescent T cells | 290 |
| Palmitate | 334 | | |

Supplementary Table 3 Number of cells for CNN model development and testing

| | Original | Training (60%) | Testing (30%) | Validation (10%) | Augmentation |
|---|---|---|---|---|---|
| Glycolysis inhibition | 2985 | 1196 | 897 | 299 | 14312 |
| OXPHOS inhibition | 2082 | 1251 | 623 | 208 | 10008 |
| Sum | 5067 | 3040 | 1520 | 507 | 24320 |

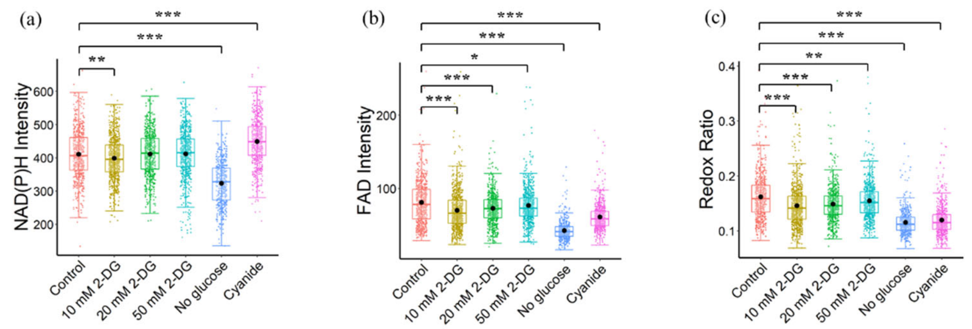


Supplementary Figure 1. Glycolysis and OXPHOS inhibition vary NAD(P)H and FAD intensity, and redox ratio. Comparison of (a) NAD(P)H intensity (b) FAD intensity (c) redox ratio (FAD/(FAD + NAD(P)H)) of cancer cells exposed to different metabolic environments. *P < 0.05, **P < 0.01, ***P < 0.001 for two-sided student test with Bonferroni correction for multiple comparisons. Substrates in each media: Control (25 mM glucose + 1 mM pyruvate), 2-DG (25 mM glucose + 1 mM pyruvate + 10/20/50 mM 2-DG), No glucose (50 mM pyruvate), Cyanide (25 mM glucose + 1 mM pyruvate + 4 mM NaCN)

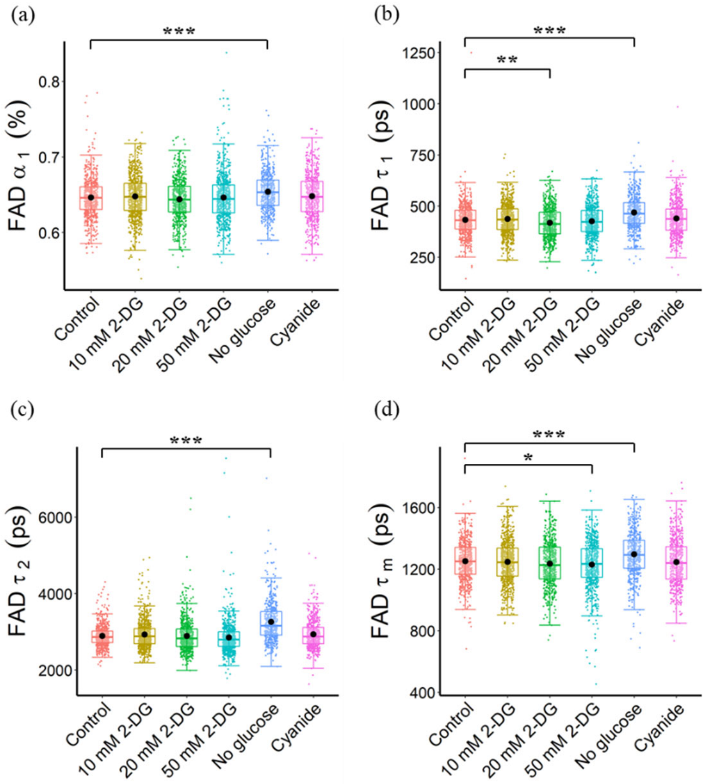


Supplementary Figure 2. Glycolysis and OXPHOS inhibition vary FAD lifetime. Comparison of (a) FAD $\alpha_1$ (b) FAD $\tau_1$ (c) FAD $\tau_2$ (d) FAD $\tau_m$ of cancer cells exposed to different metabolic environments. *P < 0.05, **P < 0.01, ***P < 0.001 for two-sided student test with Bonferroni correction for multiple comparisons. Substrates in each media: Control (25 mM glucose + 1 mM pyruvate), 2-DG (25 mM glucose + 1 mM pyruvate + 10/20/50 mM 2-DG), No glucose (50 mM pyruvate), Cyanide (25 mM glucose + 1 mM pyruvate + 4 mM NaCN)

(a)

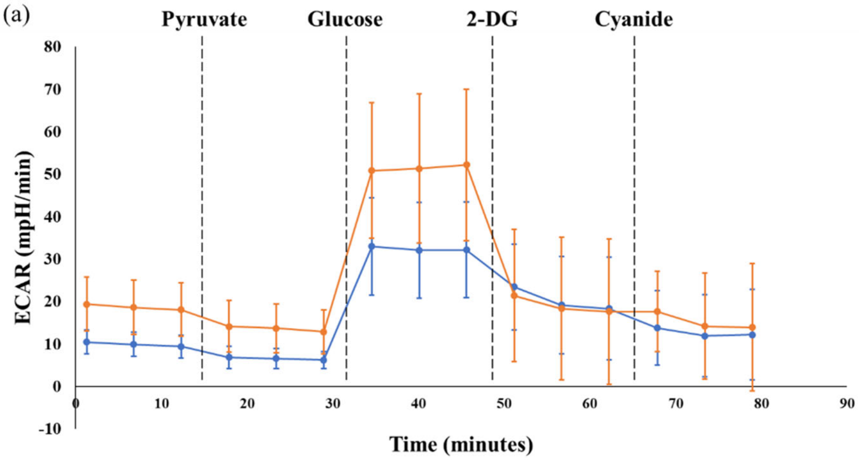


(b)

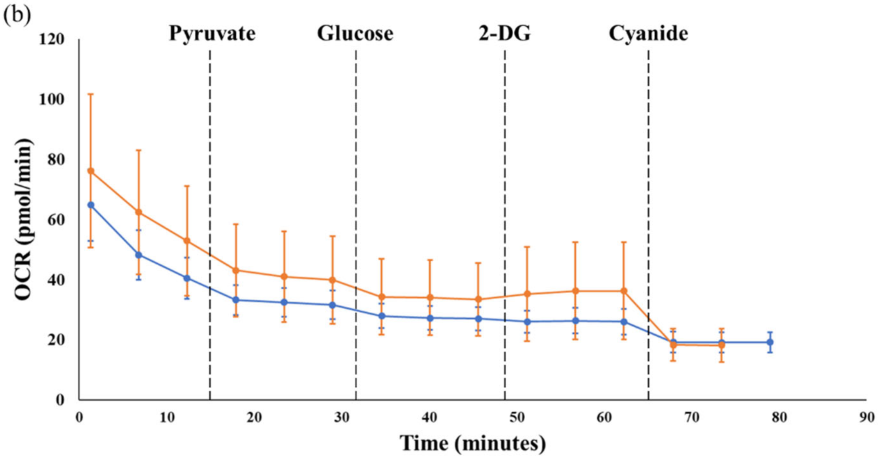


Supplementary Figure 3. Assay of mitochondrial respiration in MCF7 cells exposed to different metabolic environments. (a) Extracellular acidification rate (ECAR) and (b) oxygen consumption rate (OCR) were measured under basal conditions (no substrates) followed by the sequential addition of pyruvate (1 mM), glucose (100 mM), 2-DG (50 mM) as well as cyanide (4 mM), as indicated. Each color represents a cell group with a certain density. Orange corresponds to $10^6$ cells/ml, and blue corresponds to 5 x $10^5$ cells/ml.

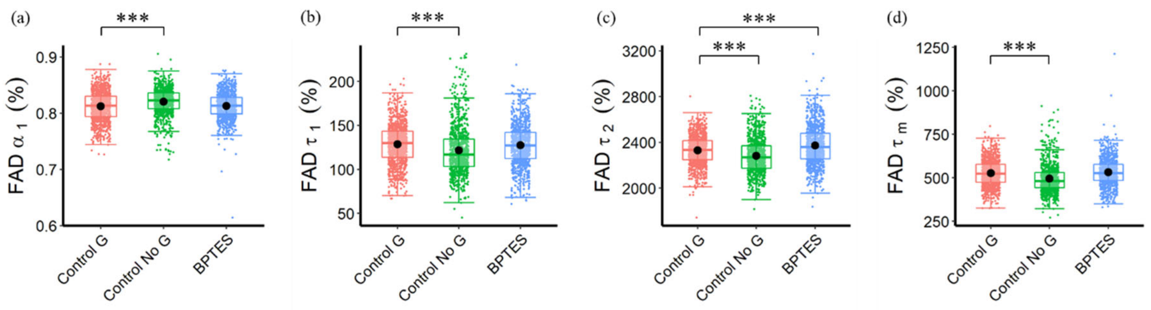


Supplementary Figure 4. Autofluorescence lifetime variations in response to glutaminolysis inhibition. (a) FAD $\alpha_1$ (b) FAD $\tau_1$ (c) FAD $\tau_2$ (e) FAD $\tau_m$ of cells in response to glutaminolysis perturbations. *P < 0.05, **P < 0.01, ***P < 0.001 for two-sided student test with Bonferroni correction for multiple comparisons. Substrates in each media: Control G (25 mM glucose + 1 mM pyruvate + 2 mM glutamine), Control No G (25 mM glucose + 1 mM pyruvate), BPTES (25 mM glucose + 1 mM pyruvate + 2 mM glutamine + 10 μm BPTES)

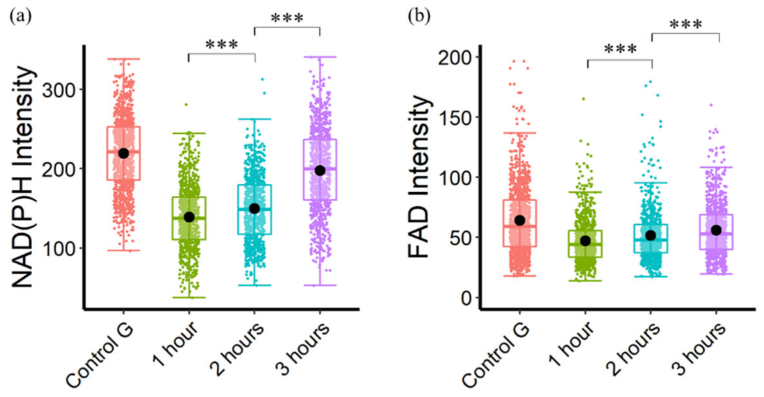


Supplementary Figure 5. Glutaminolysis effect on NAD(P)H and FAD intensity in cancer cells over the period. (a) NAD(P)H intensity (b) FAD intensity of cells exposed to only glutamine after 1 hour, 2 hours, and 3 hours. *P < 0.05, **P < 0.01, ***P < 0.001 for two-sided student test with Bonferroni correction for multiple comparisons. Substrates in each media: Control G (25 mM glucose + 1 mM pyruvate + 2 mM glutamine), 1 hour (no glucose + no pyruvate + 2 mM glutamine (1 hour)), 2 hours (no glucose + no pyruvate + 2 mM glutamine (2 hours)), 3 hours (no glucose + no pyruvate + 2 mM glutamine (3 hours)).

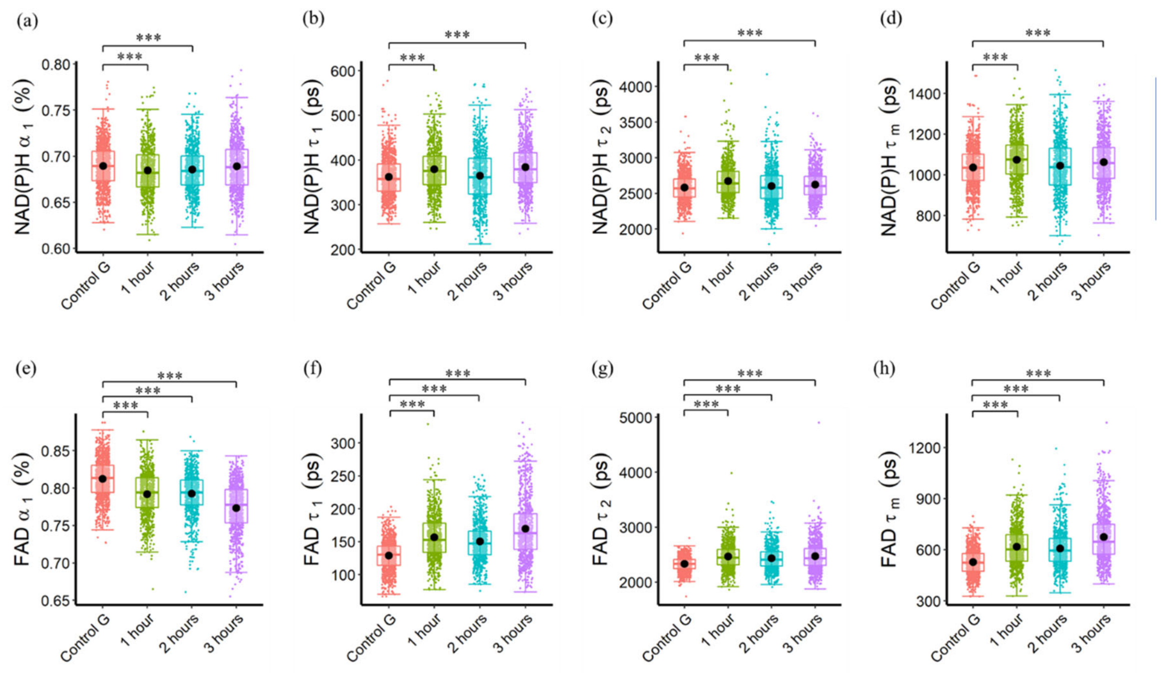


Supplementary Figure 6. Glutaminolysis effect on NAD(P)H and FAD fluorescence lifetime in cancer cells over the time period. (a) NAD(P)H $\alpha_1$ (b) NAD(P)H $\tau_1$ (c) NAD(P)H $\tau_2$ (d) NAD(P)H $\tau_m$ (e) FAD $\alpha_1$ (f) FAD $\tau_1$ (g) FAD $\tau_2$ (h) FAD $\tau_m$ of cells exposed to only glutamine after 1 hour, 2 hours, and 3 hours. *P < 0.05, **P < 0.01, ***P < 0.001 for two-sided student test with Bonferroni correction for multiple comparisons. Substrates in each media: Control G (25 mM glucose + 1 mM pyruvate + 2 mM glutamine), 1 hour (no glucose + no pyruvate + 2 mM glutamine (1 hour)), 2 hours (no glucose + no pyruvate + 2 mM glutamine (2 hours)), 3 hours (no glucose + no pyruvate + 2 mM glutamine (3 hours)).

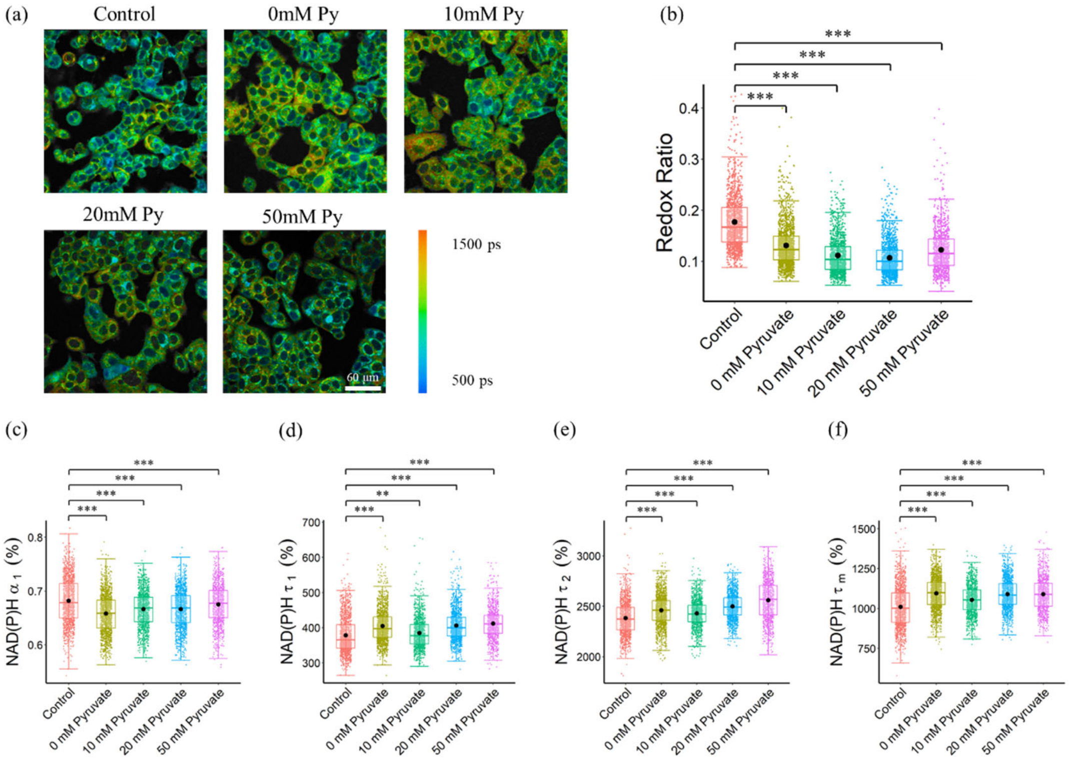


Supplementary Figure 7. NAD(P)H lifetime variations of cancer cells in different pyruvate assay groups. (a) Representative NAD(P)H $\tau_m$ images of cancer cells exposed to different pyruvate concentrations, Py, pyruvate; scale bar = 60 µm (b) Redox ratio (FAD/(FAD + NAD(P)H)) (c) NAD(P)H $\alpha_1$ (d) NAD(P)H $\tau_1$ (e) NAD(P)H $\tau_2$ (f) NAD(P)H $\tau_m$ of different pyruvate assay groups. *P < 0.05, **P < 0.01, ***P < 0.001 for two-sided student test with Bonferroni correction for multiple comparisons. Substrates in each media: Control (25 mM glucose + 1 mM pyruvate), Pyruvate (no glucose + no glutamine + 0/10/20/50 mM pyruvate).

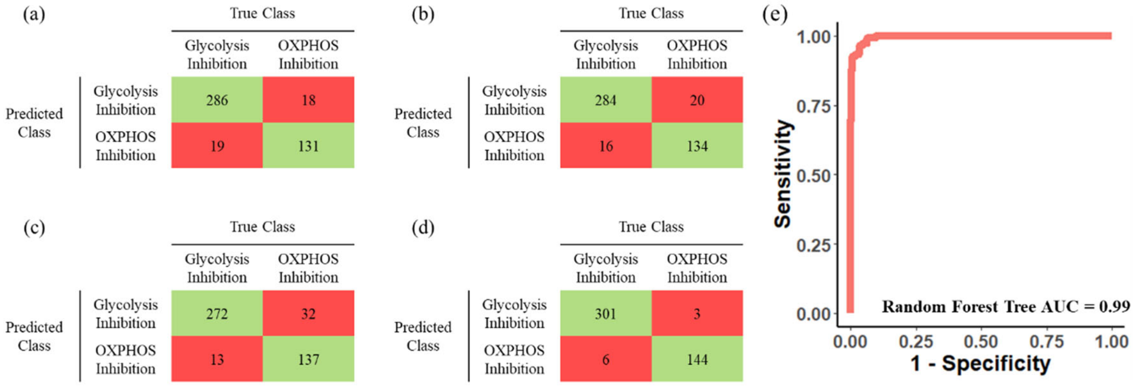


Supplementary Figure 8. Classification of cancer cell metabolic activities with autofluorescence lifetime features. Representative prediction result of (a) random forest tree (b) support vector machine (c) quadratic discriminant analysis models without feature normalization. (d) Representative prediction result of random forest tree with feature normalization by the corresponding control groups. (e) ROC curves of the test data for the random forest tree model with feature normalization by the corresponding control groups.

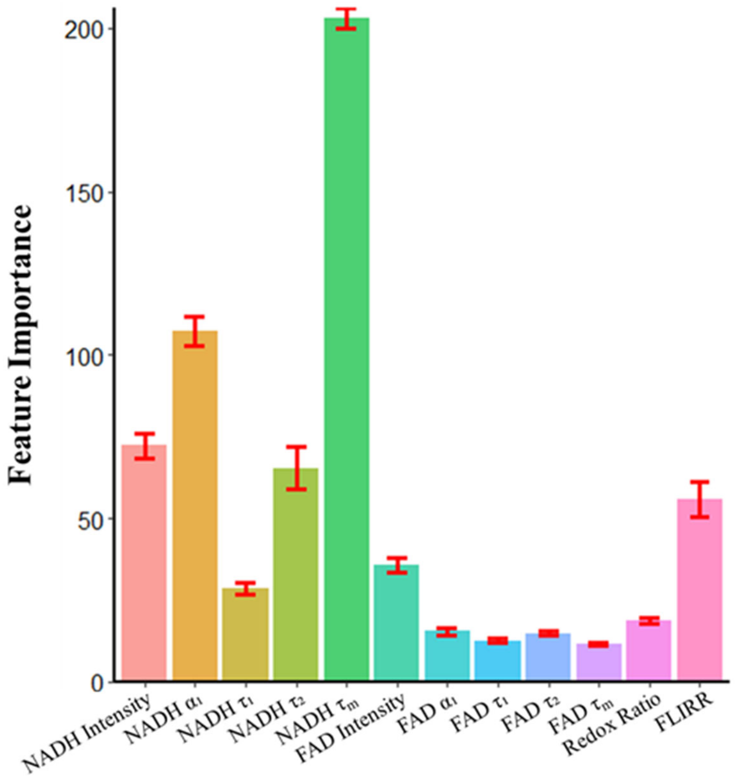


Supplementary Figure 9. Feature importance within the RFT model for classifying glycolytic versus oxidative cancer cells

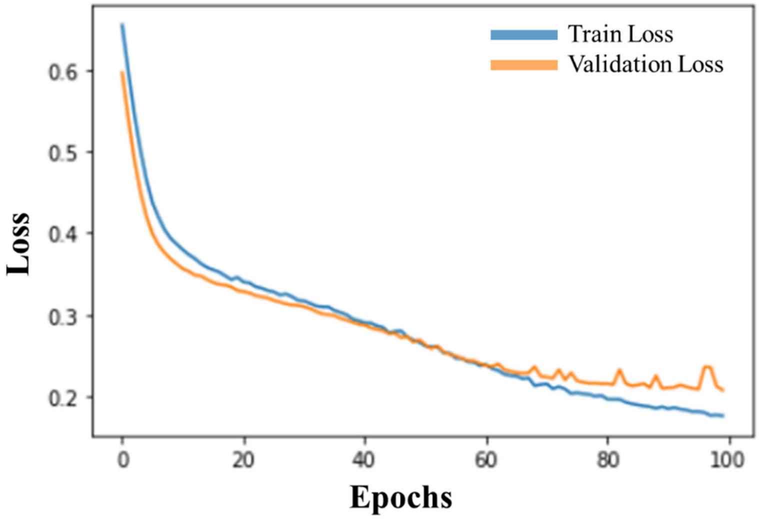


Supplementary Figure 10. Train and validation loss of CNN model (NAD(P)H $\alpha_1$ + $\tau_1$ + $\tau_2$ + $\tau_m$ + Intensity) upon the number of training epochs.

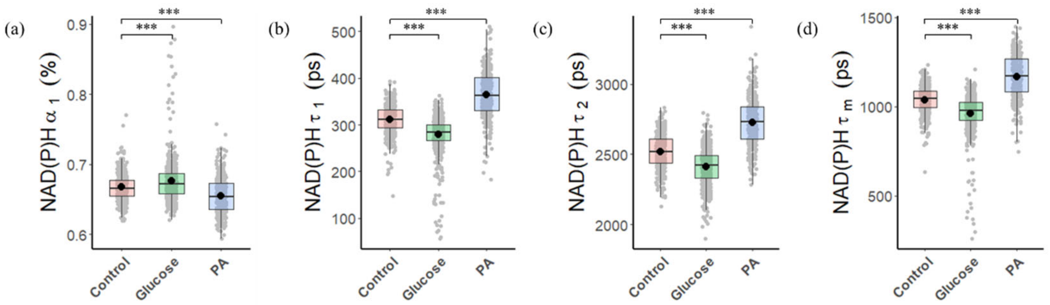


Supplementary Figure 11. Comparison of NAD(P)H lifetime components within liver cancer cells in different metabolic environments. (a) NAD(P)H $\alpha_1$ (b) NAD(P)H $\tau_1$ (c) NAD(P)H $\tau_2$ (d) NAD(P)H $\tau_m$ of cells exposed to glucose and palmitate respectively. *P < 0.05, **P < 0.01, ***P < 0.001 for two-sided student test with Bonferroni correction for multiple comparisons.

Supplementary Table 4 Prediction result of T cells with CNN model

| | Glycolysis Inhibition | OXPHOS Inhibition |
|---|---|---|
| Activated T cells | 1 (0.4%) | 262 (99.6%) |
| Quiescent T cells | 5 (1.7%) | 285 (98.3%) |
